# Excessive inflammatory and metabolic responses to acute SARS-CoV-2 infection are associated with a distinct gut microbiota composition

**DOI:** 10.1101/2021.10.26.465865

**Authors:** Werner C. Albrich, Tarini Shankar Ghosh, Sinead Ahearn-Ford, Flora Mikaeloff, Nonhlanhla Lunjani, Brian Forde, Noémie Suh, Gian-Reto Kleger, Urs Pietsch, Manuel Frischknecht, Christian Garzoni, Rossella Forlenza, Mary Horgan, Corinna Sadlier, Tommaso Rochat Negro, Jérôme Pugin, Hannah Wozniak, Andreas Cerny, Ujjwal Neogi, Paul W. O’Toole, Liam O’Mahony

## Abstract

Protection against severe acute respiratory syndrome coronavirus 2 (SARS-CoV-2) infection and associated clinical sequelae requires well-coordinated metabolic and immune responses that limit viral spread and promote recovery of damaged systems. In order to understand potential mechanisms and interactions that influence coronavirus disease 2019 (COVID-19) outcomes, we performed a multi-omics analysis on hospitalised COVID-19 patients and compared those with the most severe outcome (i.e. death) to those with severe non-fatal disease, or mild/moderate disease, that recovered. A distinct subset of 8 cytokines and 140 metabolites in sera identified those with a fatal outcome to infection. In addition, elevated levels of multiple pathobionts and lower levels of protective or anti-inflammatory microbes were observed in the faecal microbiome of those with the poorest clinical outcomes. Weighted gene correlation network analysis (WGCNA) identified modules that associated severity-associated cytokines with tryptophan metabolism, coagulation-linked fibrinopeptides, and bile acids with multiple pathobionts. In contrast, less severe clinical outcomes associated with clusters of anti-inflammatory microbes such as *Bifidobacterium* or *Ruminococcus*, short chain fatty acids (SCFAs) and IL-17A. Our study uncovered distinct mechanistic modules that link host and microbiome processes with fatal outcomes to SARS-CoV-2 infection. These features may be useful to identify at risk individuals, but also highlight a role for the microbiome in modifying hyperinflammatory responses to SARS-CoV-2 and other infectious agents.

## Introduction

Infection with SARS-CoV-2 leads to a wide variety of potential outcomes from asymptomatic responses to acute respiratory distress and death^1, 2^. While certain demographic factors such as age, male gender and comorbidities that include obesity, cardiometabolic diseases and diabetes are associated with an increased risk for more severe disease, the molecular mechanisms that underpin disease pathophysiology remain poorly understood. Indeed, we still do not know if severe outcomes are due to direct effects of viral replication within target cells, to a dysregulated host immune response to the virus, to pre-existing deficits in mechanisms of host resilience to infection, or to a combination of these factors^3,4,5^.

Initially SARS-CoV-2 infects angiotensin-converting enzyme 2 (ACE-2) expressing epithelial cells of the upper respiratory tract. If the infection remains limited to the upper respiratory tract then this is usually associated with a mild disease course and rapid recovery. If the virus is not eliminated and infection persists then other types of ACE-2 expressing cells can become infected^6^. In addition, viral-induced metabolic reprogramming and exaggerated immune responses generate a wide range of inflammatory mediators that disrupt organ homeostasis, impact host metabolism, drive a hypercoagulation state, impair epithelial barrier function and destroy host cells and tissues^7,8,9,10,11^. However, even among those who develop this cytokine storm, many can still make a full recovery, suggesting that additional factors may modulate host susceptibility to the most severe outcomes associated with COVID-19. One of these resilience factors might include the microbiome^12,13,14,15^.

Human mucosal surfaces and body cavities harbour diverse communities of commensal microbes that play essential roles in regulation of host metabolic responses, epithelial barrier function, immune education and immune regulation^16,17,18,19,20^. These effects are partially induced by activation of host pattern recognition receptors to microbial-derived danger signals, but increasingly the role for bacterial metabolites in shaping host immune function is being recognised^21,22,23^. Immunoregulatory bacterial metabolites can trigger host G protein-coupled receptors (GPCRs), aryl hydrocarbon receptors (AhRs), nuclear hormone receptors such as the farnesoid X receptor, or can directly modulate gene expression through epigenetic mechanisms. Importantly, many immunoregulatory bacterial metabolites are derived from dietary substrates (e.g. fiber), linking diet and lifestyle to protection from infection via microbial mechanisms.

In this study, our primary aim was to identify the immune-metabolic-microbial interactions and biomarkers that predict the most severe outcomes to SARS-CoV-2 infection in a well characterised cohort of patients hospitalised with COVID-19. In addition, we wished to identify clusters of patient metadata features that might provide novel mechanistic insights into the disease pathophysiology. Lastly, we wished to extend our understanding of the molecular processes within the holobiont that mediate resilience to severe biological challenges, such as viral infection.

## Results

### Systemic levels of immune mediators correlate with disease severity

While changes in circulating cytokine levels due to SARS-CoV-2 infection are already well described, the immune mediators that distinguish survivors from non-survivors in severely ill patients have not been clearly identified. To better understand the immune processes that might distinguish these patients, we measured the levels of 54 immune mediators in the earliest serum sample obtained following study enrolment after admission to the intensive care unit (ICU; severe COVID-19) or the hospital ward (mild to moderate COVID-19) from 172 hospitalised patients with PCR-confirmed SARS-CoV-2 causing COVID-19. Patient demographic details are shown in Table 1. Those with mild/moderate COVID-19 (n=42) were younger, more likely to be female, less frequently obese, required fewer medications and had fewer comorbidities compared to those with severe COVID-19 (n=130). However, there were no differences in demographics, medication use or comorbidities in those severely ill patients that survived infection (n=89), compared to those COVID-19 patients with a fatal outcome (n=41). In contrast, principal component analysis of serum immune mediators demonstrated a clear separation between patients with different COVID-19 disease outcomes (Fig. 1a). Compared to healthy volunteers (n=29), levels of 36 circulating immune mediators were significantly differed (30 higher and 6 lower) in those hospitalised with COVID-19 (Fig. 1b and Supplementary Fig. 1). Of these mediators, levels of 28 were significantly different between patients with mild/moderate COVID-19 compared to patients with severe disease (Fig. 1b). Within the severely ill group, the levels of 8 circulating immune mediators (soluble intercellular adhesion molecule-1 (sICAM-1), monocyte chemoattractant protein-1 (MCP-1), interleukin (IL)-8, macrophage-derived chemokine (MDC), interferon gamma-induced protein-10 (IP-10), IL-15, IL-1 receptor antagonist (RA) and thymic stromal lymphopoietin (TSLP)) were significantly different between those that survived and those that died (Fig. 1b and Fig. 1c).

**Table 1.** Patient Demographics

|  | Healthy Controls | Mild/Moderate | Severe - Survivors | Severe-Fatal |
| --- | --- | --- | --- | --- |
| n= | 29 | 42 | 89 | 41 |
| Age (S.D.)<sup>1,2</sup> | 45.3 (10.1) | 58.0 (15.7) | 65.0 (11.2) | 69.1 (8.7) |
| Male/Female<sup>2</sup> | 16/13 | 16/26 | 72/17 | 32/9 |
| BMI (S.D.)<sup>2</sup> | 26.9 (5.7) | 24.4 (4.1) | 28.2 (5.6) | 27.6 (5.5) |
| Obese (BMI > 30)<sup>2</sup> | 20% | 14% | 35% | 34% |
| *Medications at First Sampling Timepoint* |  |  |  |  |
| PPI<sup>2</sup> | 0% | 20% | 55% | 62% |
| Antibiotics<sup>2</sup> | 0% | 16% | 37% | 40% |
| Immunosuppressives<sup>2</sup> | 0% | 20% | 61% | 65% |
| *Pre-existing Comorbidities* |  |  |  |  |
| Hypertension |  | 34% | 49% | 45% |
| Dyslipidemia<sup>2</sup> |  | 9% | 24% | 30% |
| Diabetes |  | 17% | 22% | 25% |
| Respiratory – COPD/Asthma |  | 20% | 8% | 15% |
| Chronic Kidney Disease |  | 3% | 4% | 8% |
| Previous Neoplasia |  | 20% | 15% | 22% |

<sup>1</sup>p<0.05 Healthy Controls versus all COVID-19 Patients
<sup>2</sup>p<0.05 Mild/Moderate versus Severe COVID-19
PPI – Pantoprazole; Omeprazole
Antibiotics – Amoxicillin; Azithromycin; Sulfamethoxazole; Clarithromycin
Immunosuppressives – Dexamethasone; Methylprednisolone; Prednisolone

**Fig. 1.**
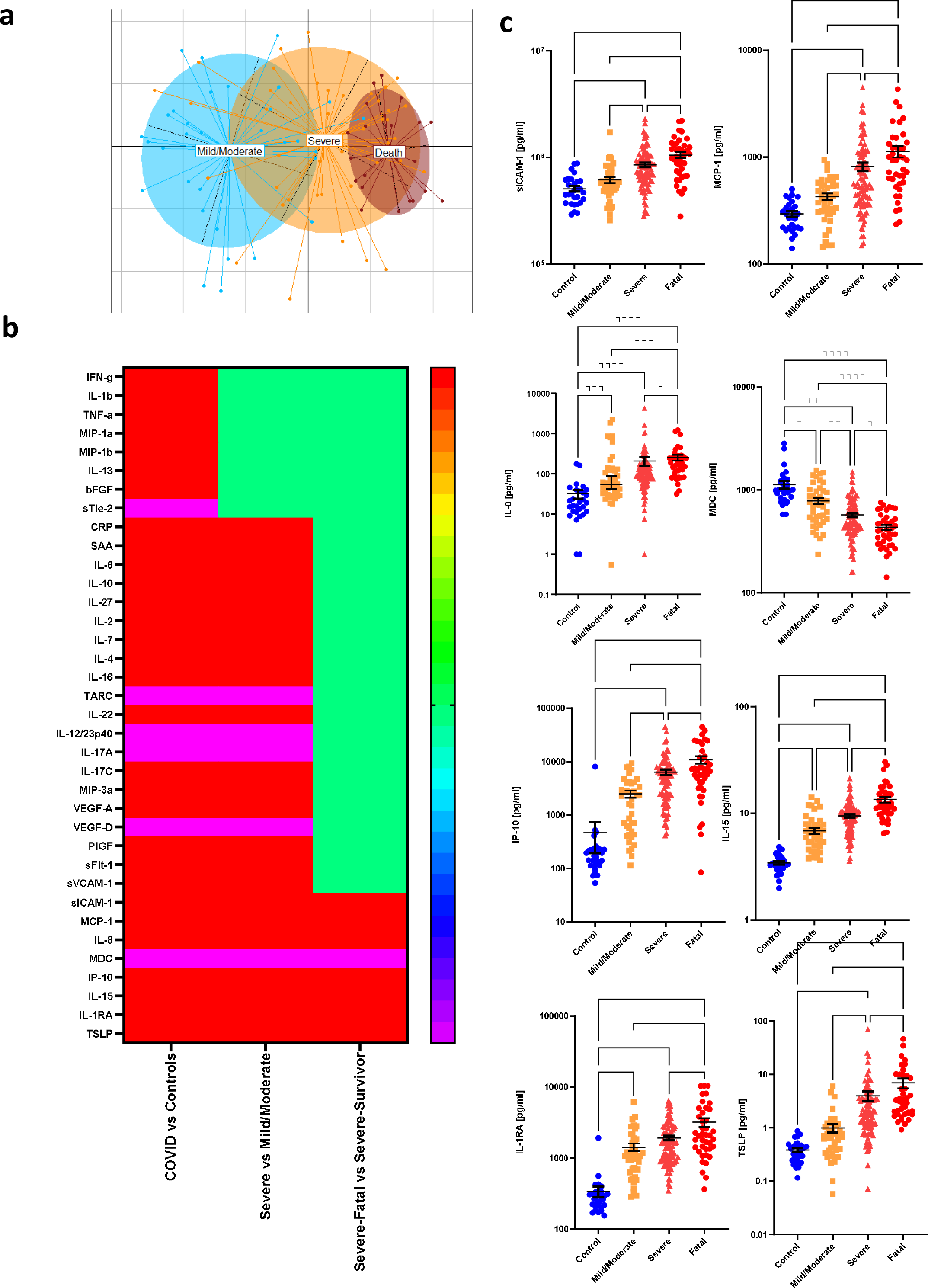
Circulating immune mediators in COVID-19 patients. a) PCA plot illustrating the differences in serum cytokine and inflammatory mediator levels in COVID-19 patients with different levels of severity. b) Heatmap illustrates the serum immune mediators that are significantly increased (red), significantly decreased (blue), or remain unchanged (green). c) Levels of the cytokines that are significantly different in patients with severe COVID-19 that survive (labelled “Severe”), compared to those with severe COVID-19 that have a fatal outcome (labelled “Fatal”). Differences between groups are calculated using the Kruskal-Wallis test and Dunn’s multiple comparison test (*p<0.05, **p<0.01, ***p<0.001, ****p<0.0001).

### Systemic metabolic responses associated with disease severity

In addition to measuring serum cytokines, we quantified and compared metabolite levels in the first serum sample obtained following study recruitment after admission to the ICU or hospital ward for COVID-19 patients with mild/moderate disease (n=25), COVID-19 patients with severe disease that survived (n=75) or COVID-19 patients with severe disease that succumbed to death (n=39). Distinct differences in circulating metabolites were evident between each of the groups (Fig. 2a and Fig. 2b). Metabolic processes were dramatically different in patients during acute SARS-CoV-2 infection, whereby levels of 377 metabolites were significantly different (adjusted p<0.05) between healthy volunteers (n=20) and those with mild/moderate COVID-19 (Fig. 2b). These differences were further exaggerated in COVID-19 patients with severe disease (583 metabolites, adjusted p<0.05), in particular those with a fatal outcome (659 metabolites, adjusted p<0.05), when compared to healthy volunteers. Within the severely ill patients, 140 metabolites distinguished those that survived versus those that died. The metabolites that contribute most to the differences between the groups included those involved in tryptophan metabolism, polyamine metabolism, histidine metabolism, lipid metabolism, bile acid metabolism and antioxidant responses such as the plasmalogens (Fig. 2c and Supplementary Fig. 2). Random forest analysis suggested a good discriminatory power for distinguishing COVID-19 disease severity or fatality based solely on a selection of circulating metabolites (Fig. 2c and Supplementary Fig. 3), underlining the robustness of these differences.

**Fig. 2.**
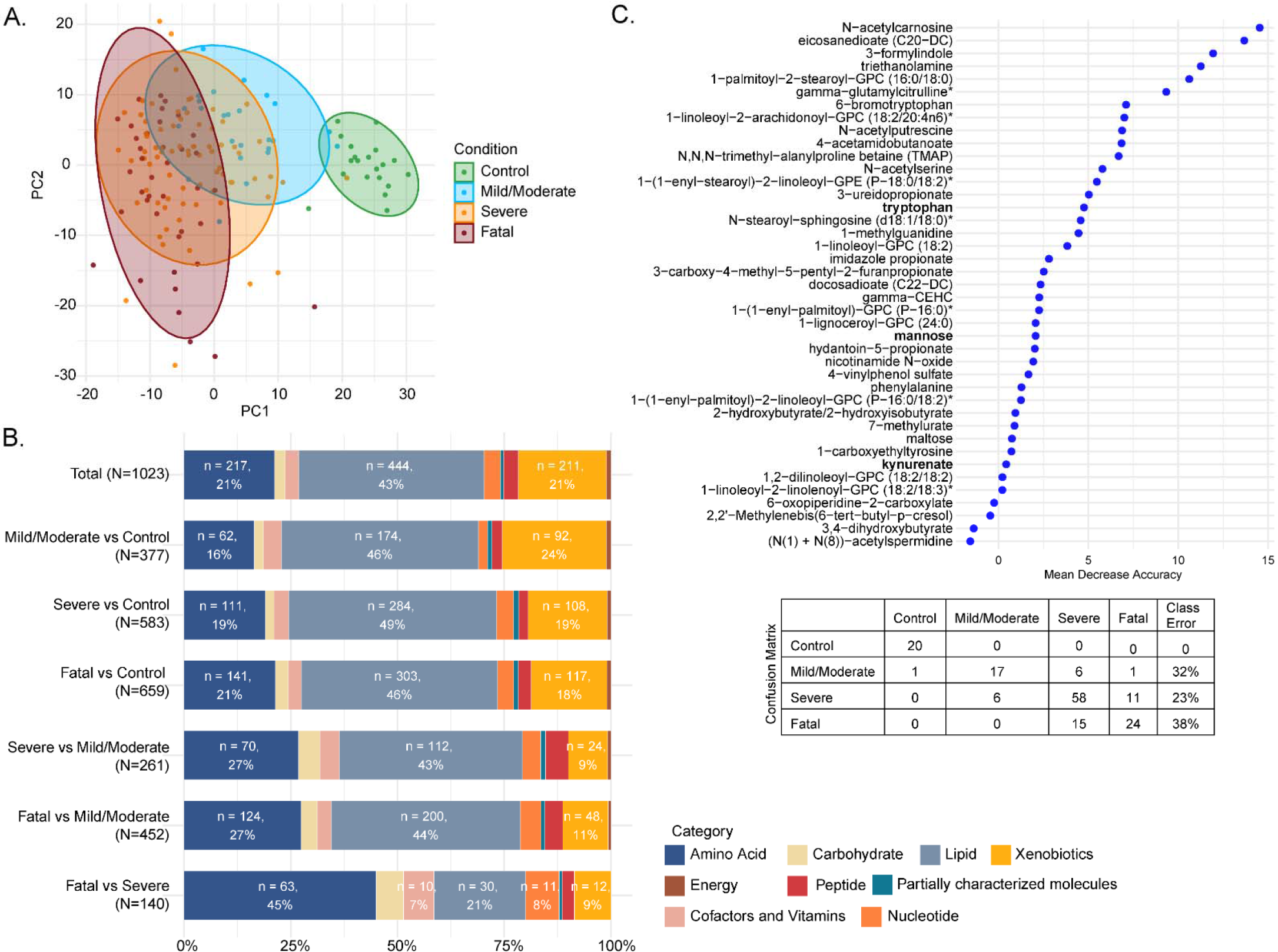
*Serum metabolites in COVID-19 patients*. a) PCA plot for the four conditions: control, mild/moderate, severe, fatal; b) Barplot representing super pathways of the significant metabolites (LIMMA, FDR<0.05) between each comparison of conditions; c) Importance plot and confusion matrix from the random forest classifier between the four conditions.

Given the substantial and significant differences in metabolite levels, we examined in more detail the most significantly impacted pathways that associated with COVID-19 severity (Fig. 3a). Interestingly, levels of sulphonated bile acids were particularly disrupted with disease severity. Host tryptophan metabolism was associated with a heavy depletion of tryptophan, with enhanced generation of kynurenate, kynurenine and quinolinate, at the expense of serotonin synthesis in COVID-19 patients (Fig. 3a and Supplementary Fig. 4a). In contrast, microbial tryptophan metabolites were present at lower levels in the serum of those with the worst outcome (Fig. 3a and Supplementary Fig. 4b). Changes in circulating microbial metabolites may be due in part to an impaired gut barrier (as indicated by increased serum SCFA levels and lower citrulline levels, Supplementary Fig. 4c and 4d), or may reflect changes in the composition or metabolism of the gut microbiome. Overall, metabolites associated with microbial metabolism (as described by Bar et al^24^) were significantly altered in those with severe disease and those with a fatal outcome (Supplementary Fig. 4e).

**Fig. 3.**
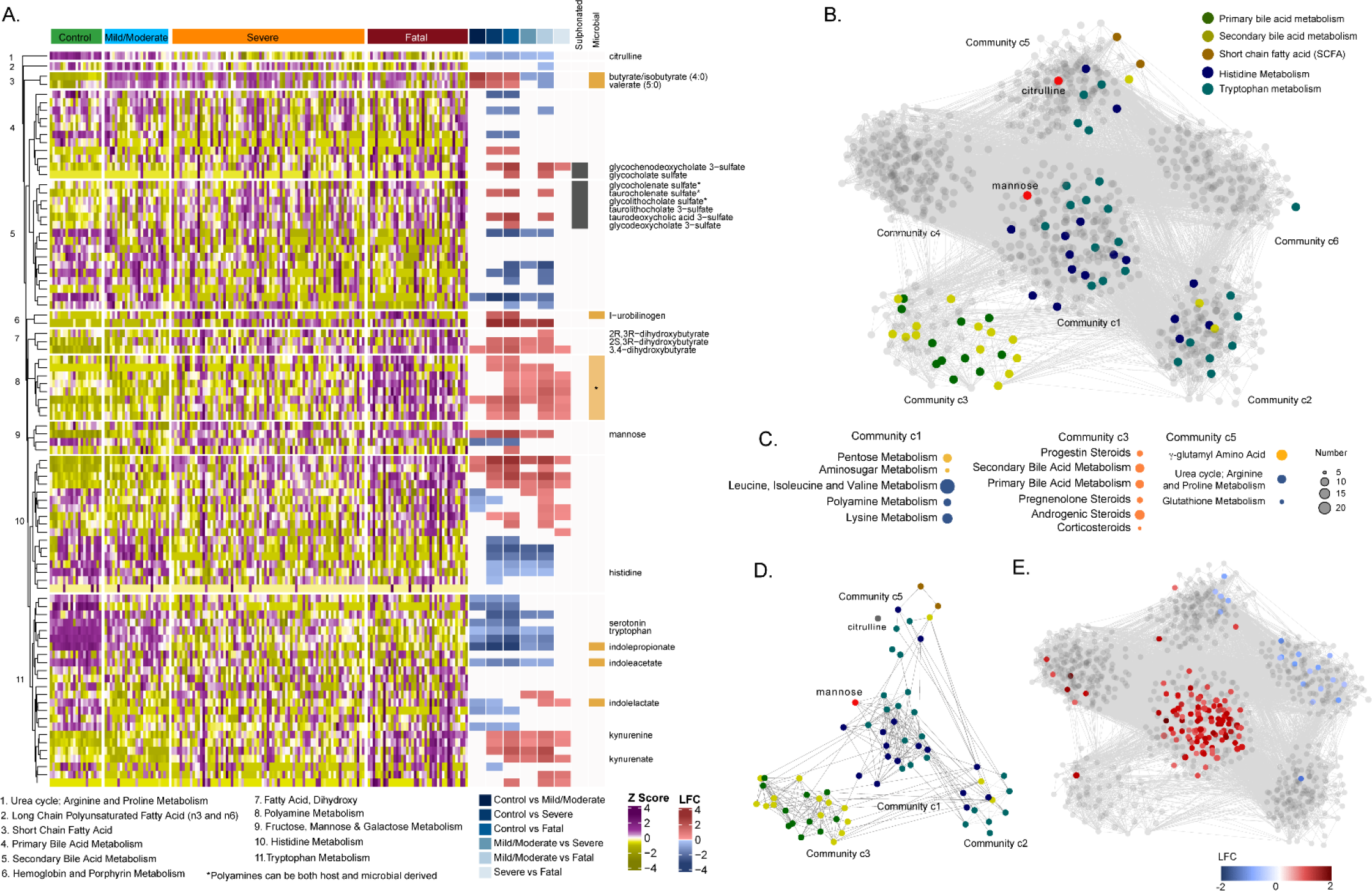
*Serum metabolites in COVID-19 patients*. a) Heatmap representing metabolites from pathways of interest, listed at the bottom of the figure, divided according to group. Log fold change (LFC) for significant pairwise comparisons (LIMMA, FDR<0.05) are included. Sulphonated bile acids and metabolites of microbial origin are indicated. b) Weighted co-expression network labelled for metabolites from pathways of interest. c) Pathway enrichment analysis using Metabolon terms for communities 1, 3 and 5 (significant terms are displayed, gseapy, FDR<0.2). d) Subset of metabolites of targeted pathways from co-expression network analysis. e) Weighted co-expression network labelled for those metabolites that were significantly different between severe COVID-19 patients that survived versus those that died (LIMMA, FDR<0.05).

**Fig. 4.**
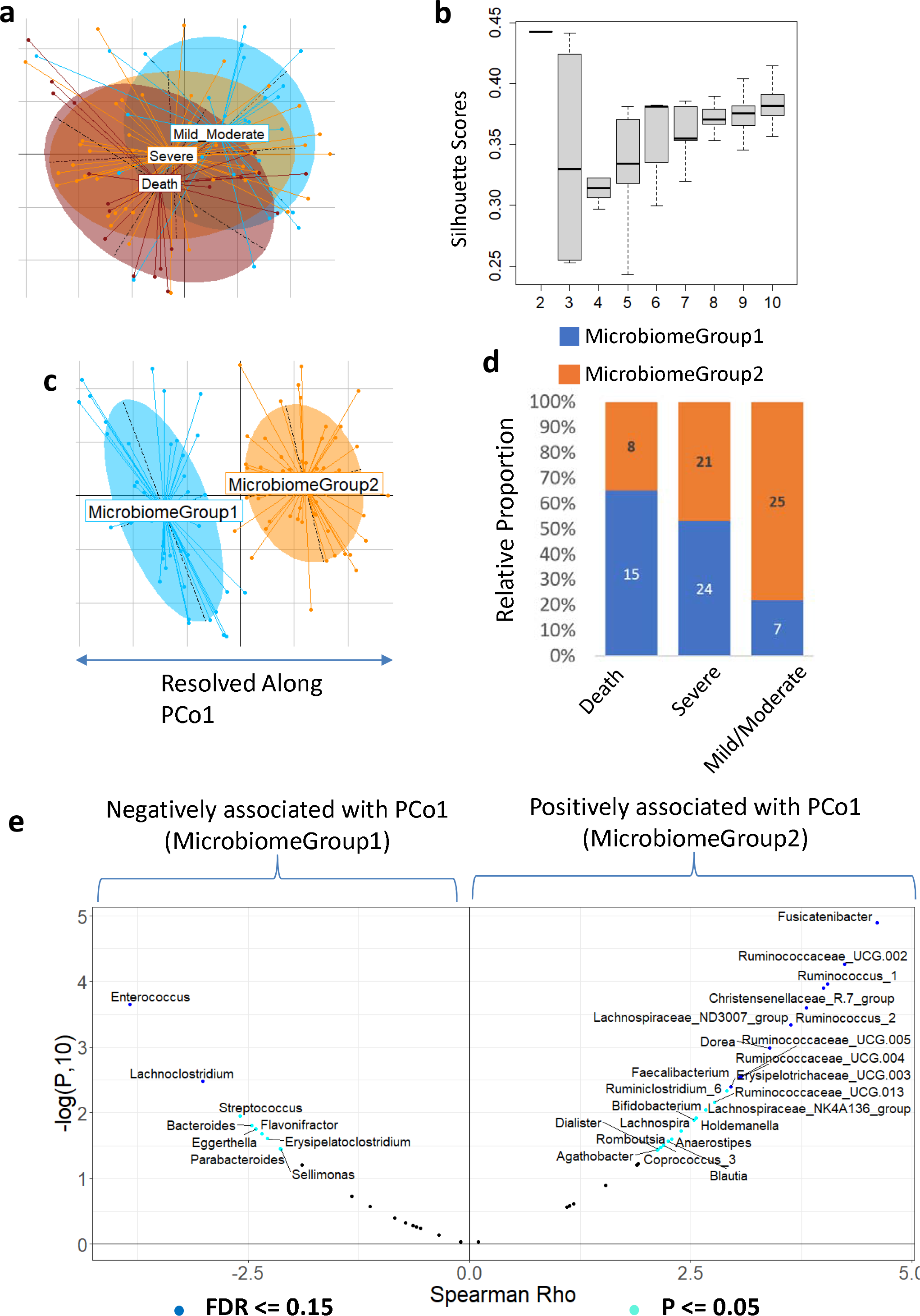
Gut microbiome composition in COVID-19 patients. (a) Principal coordinate analysis of the genus-level microbiome composition of the three outcome groups of patients obtained using the Canberra distance measure. (b) Variation of the silhouette-Scores obtained, across for cluster sizes (k), for 50 iterations of k-means clustering of the first three dominant Principal coordinates of the genus-level microbiome profiles. The principal coordinates of these two microbiome groups are demarcated in (c). The two microbiome groups exhibited distinct patterns of association with three COVID-19 disease severity outcome groups (d). Volcano plot illustrates genera showing either significant (FDR ≤ 0.15, shown in blue) or nominally significant (P ≤ 0.05, shown in cyan) associations with PCo1. The x-axis shows the estimate of the linear-regression models (direction indicating the pattern of association) and y-axis shows the -logarithm of the p-value to the base 10. The genera associating with the high-risk MicrobiomeGroup1 are on the negative axis and those associating with low-risk MicrobiomeGroup2 are on the positive axis. Only those genera showing associations with P ≤ 0.05 are shown.

Next, we performed a weighted co-expression network analysis restricted to the COVID-19 patients, to identify communities of co-abundant metabolites. Positive correlations between metabolites (Spearman, adjusted p<0.0005) were used to build the network. The analysis identified six communities (c1-c6) of highly intercorrelated metabolites based on the Leiden algorithm [2 iterations, ModularityVertexPartition, weighted network (Fig. 3b)]. Primary and secondary bile acid metabolism are contained in c3, SCFA in c5, while tryptophan and histidine metabolism are in c1, c2 and c5 (Fig. 3c). The central community (c1) with the most interconnected metabolites, central metabolites, and greatest influence on the global dynamics of the network includes mannose (Fig. 3d), which is a known inflammatory biomarker and reported to be associated with COVID-19 severity^25^. Furthermore, the metabolites that are significantly different between COVID-19 severe patients with or without a fatal outcome are primarily found within community c1 (Fig. 3e).

### Differences in the gut microbiome associate with disease severity and death

To investigate the possible involvement of the gut microbiome in these immune and metabolic changes, we profiled the microbiome by sequencing 16S rRNA gene amplicons from the first faecal samples collected following study recruitment after admission to the ICU or hospital ward for COVID-19. From the 99 hospitalised COVID-19 patients with available stool samples for 16S amplicon sequencing, 32 had mild/moderate disease, 45 had severe disease and survived, while the remaining 22 patients had severe disease with a fatal outcome. Global measures of microbiome alpha diversity were not different between clinical groups, with no significant difference detected in Shannon indices as well as in the number of detected taxa at the level of Operational Taxonomic Units (OTUs), species or genus levels between the three disease outcome groups (Supplementary Fig. 5). However, Envfit-based analysis of the Principal Coordinates revealed a significant difference in gut microbiome composition (beta diversity) between the three COVID-19 disease severity groups, irrespective of the distance measures used (Fig. 4a and Supplementary Fig. 6). We next investigated these differences in microbiome profiles in an unsupervised manner, i.e. without utilizing the disease outcome information. Using an iterative enterotyping-based approach applied on the Principal coordinates (See Methods), the microbiomes could be optimally clustered into two configurations (MicrobiomeGroup1 and MicrobiomeGroup2), resolved clearly along the first Principal Coordinate (Fig. 4b and 4c). Notably, there were significant differences in the proportions of the two distinct microbiome configurations in the clinical outcome groups (Chi-square test estimate=11.23, p-value = 0.0036, Fig. 4d). MicrobiomeGroup1 was over-represented in severe COVID-19 patients with a fatal outcome, while Microbiome Group2 was associated with those with mild/moderate symptoms. Strikingly, within the severe outcome group, individuals who were classified into the high-risk MicrobiomeGroup1 had significantly higher levels of cytokines associated with both fatality and severity (P = 0.02; Mann-Whitney Test), with higher (albeit not statistically significant) levels of cytokines associated only with disease severity (P = 0.12, Mann-Whitney Test) (Supplementary Fig. 7).

We next investigated the genus-level composition differences across the two microbiome configurations by performing ordinary-least square (OLS)-based regression analysis to measure the association between abundance of microbial genera and the PCo1 axis values after adjusting for confounders. A total of 9 genera showed significant associations with PCo1 with FDR ≤ 0.15 (Benjamini-Hochberg corrected), even after confounder adjustment. While two genus-level groups (*Enterococcus* and an unclassified member of the *Enterococcaceae*) were associated negatively with PCo1 (high relative abundance in the high risk MicrobiomeGroup1), the other 7 (comprising *Christensenellaceae*-R7, *Dorea*, *Fusicatenibacter* and multiple *Ruminococcus* species) showed the opposite trend (Fig. 4e). Relaxing the thresholds identified 19 more genera that showed nominally significant association with PCo1 (P ≤ 0.05). The high risk MicrobiomeGroup1 was characterized by higher levels of multiple pathobionts (as operationally defined in our previous work^26, 27^) including *Enterococcus*, *Eggerthella*, *Lachnoclostridium, Erysipelatoclostridium, Streptococcus, Flavonifractor* and lower levels of multiple taxa known to be associated with anti-inflammatory or protective immune responses (including *Faecalibacterium, Agathobacter, Dorea, Coprococcus, Lachnospiraceae, Christensenellaceae*) (Fig. 4e). Many of the observed differences in the microbiome were significantly associated with changes in levels of circulating immune mediators (Supplementary Fig. 8).

### Immune-metabolite-microbiome modules correlate with COVID-19 disease outcomes

Correlation network analysis is a powerful tool for revealing associations of diverse features within patient datasets. Feature-association networks were computed using the Weighted gene correlation network analysis (WGCNA) approach (see Methods) performed on 1,469 features (54 cytokines, 1,146 metabolites and 269 microbial genera) using signed Spearman correlations with a soft-power threshold of 7 (Supplementary Figure 9a) from the 70 hospitalised COVID-19 patients with complete data for all three data layers. A total of 14 modules (annotated as different colors) were identified, 5 of which had a significant association with disease outcome (Benjamini-Hochberg FDR ≤ 0.05) and 2 modules showed nominal associations (P ≤ 0.05 and FDR ≤ 0.1) (Supplementary Fig. 9b-c). The module (annotated as ‘turquoise’) that showed significant positive association with disease severity and death contained most of the severity associated cytokines (as identified in Fig. 1), metabolites (Supplementary Fig. 3) and microbial genera identified above (Fig. 4e), combined with kynurenine associated metabolism products and coagulation linked fibrinopeptides (Fig. 5a). Two modules (annotated as ‘brown’ and ‘tan’) were nominally positively associated with a poor outcome. Of these, the brown module contained a triad of pathobionts linked to urobilinogen (Supplementary Fig. 10a), while the tan module was enriched for sulfonated bile acids (Supplementary Fig. 10b). In contrast, 4 modules, annotated as ‘red’, ‘blue’, ‘black’ and ‘yellow’, were significantly negatively associated with COVID-19 severity and death. The first module (red) contained the anti-inflammatory *Ruminococcus*_2 clade, linked with tryptophan, alanine and the SCFAs butyrate/isobutyrate and valerate (Fig. 5b; Supplementary Fig. 11). The second module (blue) that negatively associated with disease severity contains a cluster of beneficial microbial taxa (including *Bifidobacterium*), bilirubin degradation products, TARC and IL-17A (Supplementary Fig. 12). The third module (black) exclusively contains metabolites, in particular fatty acid derivatives (Supplementary Fig. 13), while the final significant module (yellow) contains *Roseburia, Fusicatenibacter, Romboutsia* linked with sphingomyelin and carnitine-derived products (Supplementary Fig. 14).

**Fig. 5.**
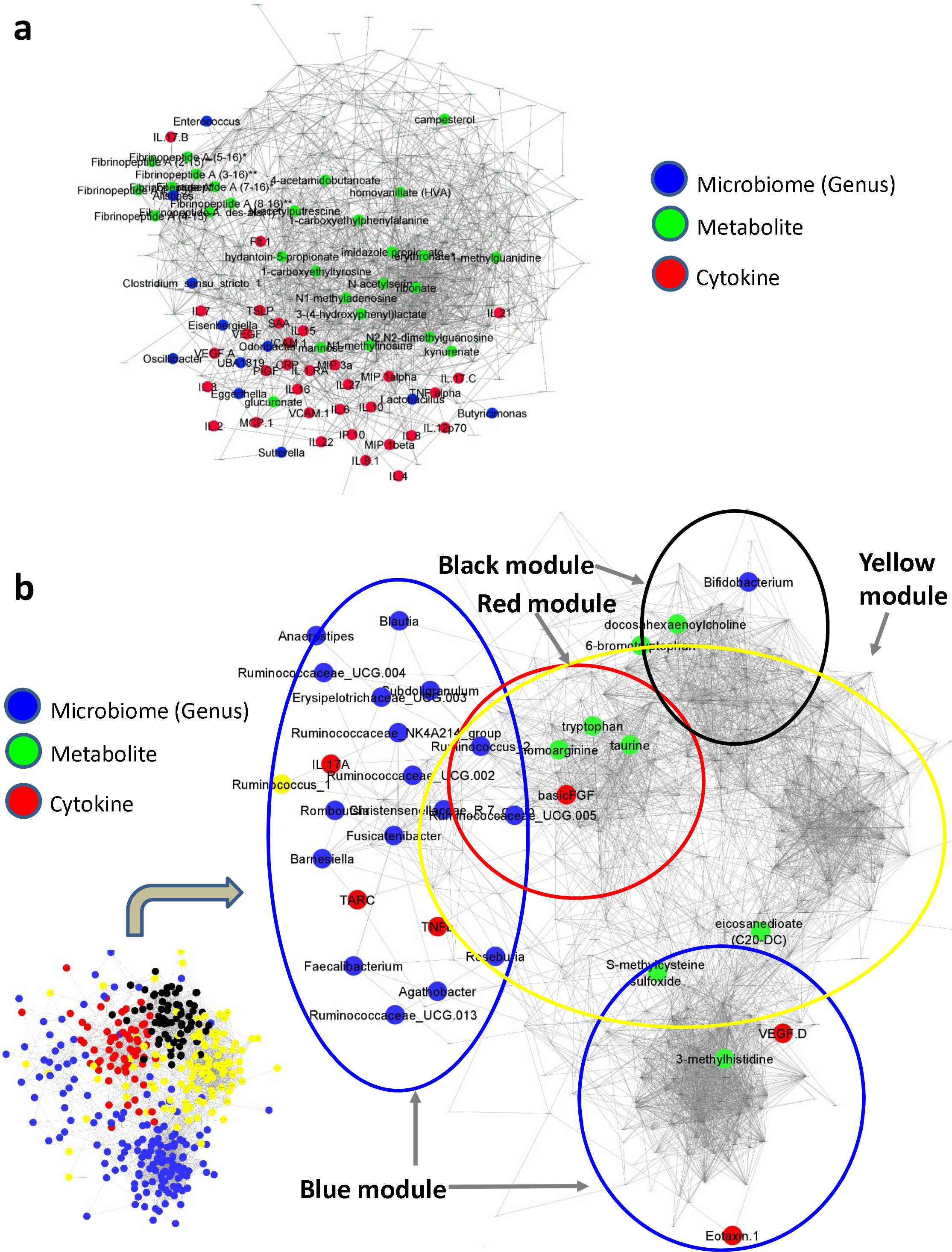
Modules that positively correlate with severe and fatal COVID-19. Feature-to-feature positive association networks obtained using the ccrepe approach (Spearman correlations, 1000 iterations) for modules (or Module groups) that show (a) significantly positive (‘turquoise’) and (b) significantly negative (‘red’, ‘blue’, ‘yellow’, and ‘black’) associations with severe and fatal COVID-19. In (b) given the presence of features from four different modules, the location of the features belonging to the different modules are indicated in the smaller network representation in the lower left-hand corner. Microbiome, cytokine and metabolite features that are associated with severity and death are highlighted in different colours.

## Discussion

Despite the substantial literature published on SARS-CoV-2, the molecular mechanisms underpinning positive versus negative clinical outcomes remain poorly defined. In this study, we examined the differences in circulating inflammatory markers and metabolites in sera, and the composition of the gut microbiota, in a large group of hospitalised patients with COVID-19. We have identified several potential regulatory nodes whereby integrated immune, metabolic and microbiome processes contribute to susceptibility or resilience to SARS-CoV-2 infection associated damage.

Our identification of circulating inflammatory mediators that associate with COVID-19 disease severity such as CRP and IL-6 are consistent with previous reports and support the hypothesis that an overly aggressive immune response contributes to immunopathology and severity^28, 29^. In addition to severity associated factors, we have identified a subset of eight cytokines that are further dysregulated in severe patients with a fatal outcome. Higher levels of IP-10 and IL-15 indicate greater activation of a T helper 1 (Th1)-associated innate anti-viral response, while a significant reduction in MDC levels may reflect the inhibitory effect of a Th1 environment on Th2 cytokines such as MDC. We were particularly interested in TSLP as this cytokine is an epithelial cell-derived alarmin, which is released by injured stromal cells to recruit and activate innate immune cells, and its blockade is currently being investigated in asthma clinical studies^30,31,32^. In combination with the chemokines MCP-1 and IL-8, and sICAM-1 (which modulates leukocyte adhesion and migration across endothelial cells), elevated TSLP levels indicate a greater amount of epithelial tissue damage and inflammatory cell recruitment to the damaged sites in patients who do not recover from SARS-CoV-2 infection. As SARS-CoV-2 is a lytic virus, it is possible that viral replication in epithelial cells may directly drive TSLP levels in sera, although indirect effects on epithelial cells within the respiratory tract or gut might also induce TSLP release. Importantly, TSLP levels were previously shown to be elevated in patients with long COVID, suggesting that long term impacts of SARS-CoV-2 on epithelial cells should be examined in more detail, potentially guiding future therapeutic interventions^33^.

Significant metabolic reprogramming and compensatory responses are evident in COVID-19 patients with severe disease and particularly in those with a fatal outcome. Decreased serum levels of plasmalogens suggest a significant level of systemic oxidative stress as these sacrificial phospholipids are preferentially oxidised to protect more vulnerable membrane lipids such as polyunsaturated fatty acids^34^. Altered tryptophan metabolism was particularly interesting to observe as the profound shutdown in serotonin production coupled with accumulation of quinolinic acid indicated a shift from production of neuroprotective compounds to production of neurotoxic compounds, which might be clinically relevant^35^. An imbalance between host and microbial tryptophan metabolism was also evident as serum kynurenine levels increased, while products of bacterial tryptophan metabolism such as indoleacetic acid were significantly decreased in those with severe and fatal disease^36^. These are important AhR ligands that can contribute to immune regulatory responses, can drive an “exhaustion” phenotype in immune effector cells, and are important for maintenance of the gut epithelial barrier by induction of IL-22^37, 38^. Other significantly different metabolites such as the polyamines putrescine and spermidine play important roles in protecting against inflammatory responses within the airways^39^. In addition, changes in secondary bile acid serum levels indicate significant disruption of microbial metabolism and/or changes in the gut barrier.

Secondary bile acids significantly impact regulatory and effector immune responses, which may be relevant for the development of severe COVID-19^40, 41^. Increased levels of sulfonated bile acids in serum also indicates significant disruption of bile acid metabolism in severely ill COVID-19 patients as sulfonation is an important detoxification mechanism that prevents reabsorption of bile acids from the gut and promotes their elimination in faeces^42^.

We identified a high-risk gut microbiome configuration associated with an inflamed host phenotype and increased risk of the worst disease outcomes. Several pathobionts including *Enterococcus* were enriched in severe disease, while well described immune regulatory microbes such as *Bifidobacterium* and *Ruminococcus* were enriched in those who survived^43, 44^. Similar microbiome configurations have been described in other settings such as increasing age, whereby a decrease of the core protective microbiome accompanied by an increase of pathobionts was observed^45^. In addition, acquisition of this subset of disease-associated taxa have been shown to shift the metabolic state to a disease-like state^27^. These changes in the microbiome may have happened gradually over time and could potentially make the host less resilient to SARS-CoV-2 infection.

The hyper-inflammatory state observed in COVID-19 patients with a fatal outcome implies a failure in the negative feedback mechanisms that should restrain the devastating overproduction of inflammatory cytokines and soluble mediators, which lead to multiorgan failure. Our integrated analysis of microbiome features, cytokines and metabolites suggests that important microbial-derived immunoregulatory processes that contribute to negative feedback mechanisms may be lacking in those with the most severe outcomes to SARS-CoV-2 infection. Alternatively, increased levels of proinflammatory pathobionts may drive excessive proinflammatory responses that cannot be contained by the regular feedback mechanisms. While further studies will be required to determine causal interactions, this study supports the hypothesis that successful responses to infectious agents such as SARS-CoV-2 involve the gut microbiome mediated by effects on metabolism and host inflammatory processes.

## Methods

### Study Cohort

We performed an investigator-initiated, prospective multicentre cohort study of adult (≥ 18 years) patients who were admitted with Severe Acute Respiratory Syndrome Coronavirus 2 (SARS-CoV-2) to four different hospitals in Switzerland and Ireland. Infection was confirmed by SARS-CoV-2 polymerase-chain reaction (PCR) from an upper or lower respiratory specimen. Exclusion criteria included COVID-19 diagnosis after discharge from the ICU. Recruitment started in August 2020 and in total we recruited 172 hospitalised patients from St. Gallen, Switzerland (n=37), Geneva, Switzerland (n=50), Ticino, Switzerland (n=77) and Cork, Ireland (n=8). All patients or patient representatives signed a patient informed consent. The study was approved by local ethics committees (EKOS 20/058 for the three Swiss sites and The Clinical Research Ethics Committee of the Cork Teaching Hospitals for Cork University Hospital). Patients were enrolled typically within 24-48 h after admission to the intensive care unit (ICU) or a hospital ward. Baseline characteristics, underlying comorbidities and medication use at the time of sampling were collected and are summarised in Table 1. All medical procedures and treatments were left at the discretion of the treating physicians but documented in the database such as complications during ICU stay and outcomes until hospital discharge. Patients were categorised to have mild disease when there were no radiographic indications of pneumonia and moderate disease if pneumonia with fever and respiratory tract symptoms were present. Severe disease was defined as a respiratory rate ≥30 breaths per minute, oxygen saturation ≤93% when breathing ambient air or PaO2/FiO2 ≤300mm Hg, or anyone that required mechanical ventilation. Only those that died during their hospital stay were recorded as a SARS-CoV-2-related death in this study. Serum and faecal samples were collected as soon as possible following enrolment into the study and immediately stored frozen at -80C at the clinical site.

### Cytokine Analysis

We examined the levels of 54 cytokines and growth factors (using MSD multiplex kits according to manufacturer’s instructions) in the serum of 172 hospitalised COVID-19 patients. Serum from patients was typically obtained within 24 hours after study enrolment. Sera obtained prior to the pandemic from 29 healthy volunteers were analysed in parallel. The mediators measured included IL-1α, IL-1β, IL-1RA, IL-2, IL-3, IL-4, IL-5, IL-6, IL-7, IL-8, IL-9, IL-10, IL-12/23p40, IL-12p70, IL-13, IL-15, IL-16, IL-17A, IL-17A/F, IL-17B, IL-17C, IL-21, IL-22, IL-23, IL-27, IL-31, TNF-α, TNF-β, IFN-γ, IP-10, MIP-1α, MIP-1β, MIP-3α, MCP-1, MCP-4, Eotaxin, Eotaxin-3, TARC, MDC, TSLP, CRP, SAA, VEGF-A, VEGF-C, VEGF-D, sTie-2, Flt-1, sICAM-1, sVCAM-1, bFGF, PIGF and GM-CSF.

### Metabolomics

Untargeted metabolomics on patient sera was performed by Metabolon^TM^ using the HD4 platform. Briefly, all methods utilized a Waters ACQUITY ultra-performance liquid chromatography (UPLC) and a Thermo Scientific Q-Exactive high resolution/accurate mass spectrometer interfaced with a heated electrospray ionization (HESI-II) source and Orbitrap mass analyzer operated at 35,000 mass resolution. The sample extract was dried then reconstituted in solvents compatible to each of the four methods. One aliquot was analyzed using acidic positive ion conditions, chromatographically optimized for more hydrophilic compounds. Another aliquot was also analyzed using acidic positive ion conditions, however it was chromatographically optimized for more hydrophobic compounds. Another aliquot was analyzed using basic negative ion optimized conditions using a separate dedicated C18 column. The fourth aliquot was analyzed via negative ionization following elution from a HILIC column (Waters UPLC BEH Amide 2.1x150 mm, 1.7 µm) using a gradient consisting of water and acetonitrile with 10mM Ammonium Formate, pH 10.8. The MS analysis alternated between MS and data-dependent MSn scans using dynamic exclusion. The scan range varied slighted between methods but covered 70-1000 m/z.

### 16S sequencing

Fecal samples were obtained as soon as possible following hospitalisation. Total community DNA was extracted from fecal samples by a combined Repeat Bead Beating - Qiagen DNA extraction method, and the V3 dash V4 region of the 16S gene was amplified and sequenced as previously described^46^. The uniquely barcoded amplicons were sequenced on an Illumina MiSeq platform (Illumina, California, USA) utilizing 2×300 bp chemistry.

### Bioinformatic analysis

From the Log2 transformed metabolomics data obtained from Metabolon, any metabolite with no variance among samples was removed. Pairwise differential abundance analysis was performed between conditions using R package LIMMA. Benjamini-Hochberg correction (BH) was applied for each comparison. R packages Boruta was applied for feature and tree number selection before random forest analysis. Random forest classifiers were built with the most important features, 1000 trees, mtry of 1 and 10-fold cross-validation using R packages caret and randomForest. They were evaluated using confusion matrices/roc curves. For association analysis, significant positive correlations (Spearman, FDR<0.0005) were extracted and used to build the network using python igraph (https://igraph.org/python/). The strength of the connections and relevance of the network was evaluated by plotting distribution of correlation coefficients and comparison of the network to a random network with similar dimensions. Community detection was performed using the Leiden algorithm from the python module leidenalg (https://leidenalg.readthedocs.io/en/stable/index.html). For each community large enough (N>30), metabolite set enrichment analysis (MSEA) was performed. For metabolite set enrichment analysis (MSEA), all Metabolon^TM^ terms were extracted with their corresponding metabolites as reference. Python 3 gseapy package was used to perform a hypergeometric test between list of significant metabolites and reference. Importance plots, dot plots, bar plots, pca plots were produced with R package ggplot2. Heatmaps were designed with the R package ComplexeHeatmap. Networks were represented using Cytoscape 3.6.1 and metabolites of interest highlighted.

For the microbiome analysis, the raw Illumina reads obtained for each sample were quality-filtered using the trimmomatic program, using the default parameters^47^. The quality filtered reads were then taxonomically classified using both DADA2^48^ (for read-level genus classification and identification of amplicon sequence variants or ASVs within each sample) and Spingo^49^ (for species level classification). Amplicon Sequence Variants obtained using DADA2 for all the samples were then further merged by performing into Operational Taxonomic Units (OTUs) using the denovo-sequence-based clustering using the qiime^50^ package.

Subsequent downstream analyses of the taxonomic profiles (at all three levels, namely genus, species and OTU) as well as integrated analysis of taxonomic profiles with cytokine profiles and the metabolome were performed using various modules/packages of the R programming interface (v 4.0.3; R Core Team 2020). Estimates of alpha diversity were computed using the diversity function of the vegan package of R. Principal Coordinate Analyses (PCoA) were computed using the ade4 package. The envfit function of the vegan package was used to perform the envfit-based analysis using the three top Principal Coordinates. Enterotyping of the gut microbiome profiles was performed as described in a previous study from our group^51^. Two group comparison of microbiome abundances were performed using the Mann-Whitney tests (using the wilcox.test function of R stats package. For more than two-group comparisons, pairwise comparisons within groups were computed using Mann-Whitney tests. The p-values were corrected using Benjamini-Hochberg FDR correction (p.adjust function of the stats package). Ordinary Least Squares (OLS) regression after adjusting for confounders were performed using the glm function of the stats package.

Correlation analysis of associations amongst features in three data layers (genus-level microbiome, metabolome and cytokine profiles) were performed using the Weighted Gene-Coexpression Network Analysis (WGCNA)^52^. While originally devised for computing gene co-expression networks, WGCNA is now being used in studies to integrate data from multiple OMICs layers^53, 54^. In this study, the WGCNA was performed using an optimal soft-power threshold of 7 for scale-free topology. Using hierarchical clustering and topology overlap measures (TOM), we identified that the features from the three data layers could be optimally grouped into 14 modules, which were then investigated for association with disease symptoms using OLS models. The association networks within each module were then computed using the ReBoot approach as implemented in the ccrepe workflow^55^.

## Acknowledgements

The authors are supported in part by Science Foundation Ireland (SFI) in the form of a research center grant to APC Microbiome Ireland (12/RC/2273_P2) and two COVID-RRC awards (20/COV/0158 (LOM) and 20/COV/0125 (PWOT)). Work in UN’s laboratory is supported by the Swedish Research Council grants (2017-01330 and 2018-06156). In addition, funding from an Intramural grant, Cantonal Hospital St. Gallen, Fondazione Leonardo and Fondazione Metis Mantegazza supported this study.

For their valuable contributions to this study, we would like to thank Susan Rafferty-McArdle, Mary Crowley, Manasi Nadkarni, John MacSharry, Liam Fanning, Brian McSharry (University College Cork); Ines Thiele (University College Galway); Christian Kahlert, Miodrag Filipovic, Cornelia Knapp, Susanne Nigg, Thomas Egger, Andrea Blöchlinger, Tia Wisser, Melanie Gätzi and Patrick Münger (Cantonal Hospital St. Gallen); Kenza Bouras, Aurélie Perret and Philippe Montillier (Geneva University Hospitals and the University of Geneva Faculty of Medicine); Maurizia Bissig-Canevascini and Claudia Di Bartolomeo (Fondazione Epatocentro Ticino).

## Supplementary Figure Legends

**Supplementary Fig. 1.**
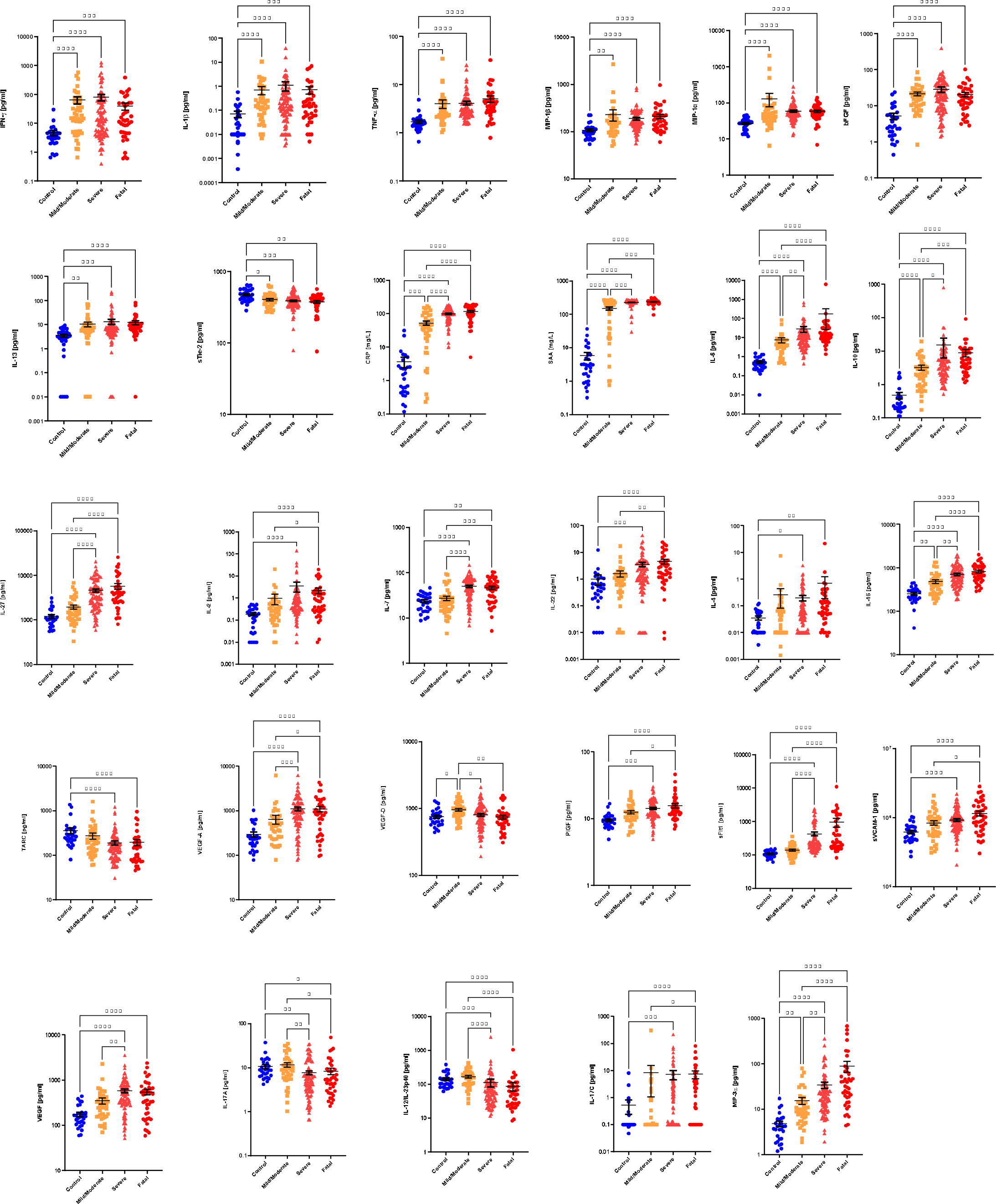
Serum cytokine levels. Differences between groups are calculated using the Kruskal-Wallis test and Dunn’s multiple comparison test (*p<0.05, **p<0.01, ***p<0.001, ****p<0.0001).

**Supplementary Fig. 2.**
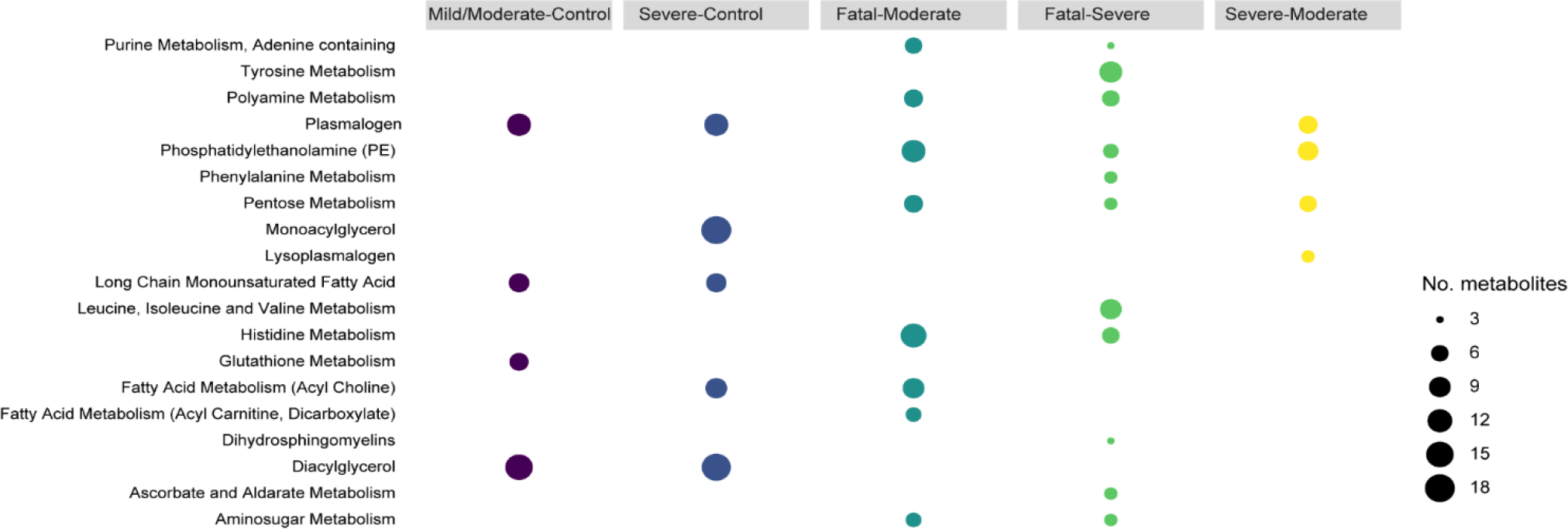
Metabolite set enrichment analysis. Using Metabolon terms (gseapy, FDR < 0.2), bubble size represents the number of metabolites found in each pathway. Color is specific to each comparison.

**Supplementary Fig. 3.**
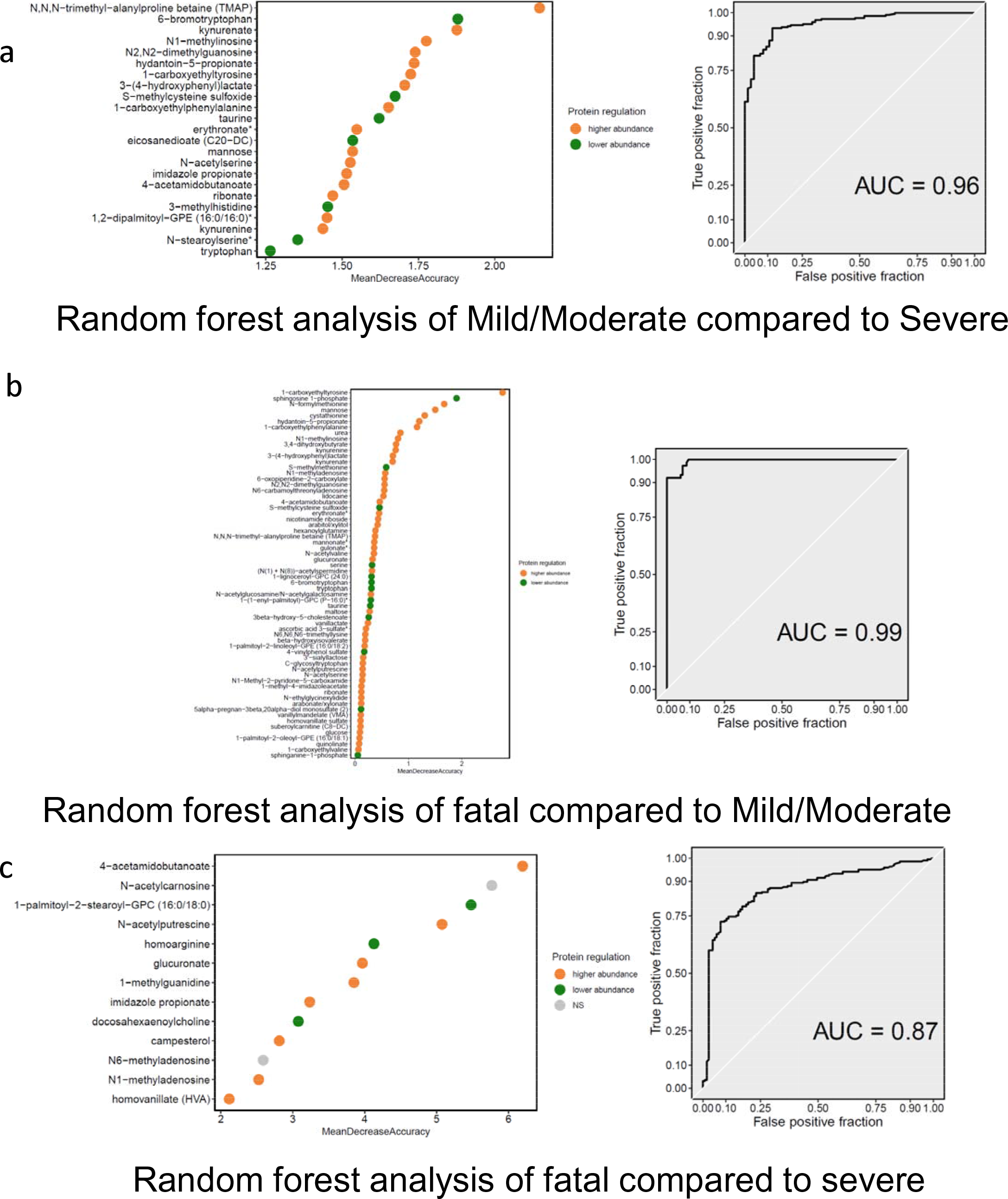
Random forest analysis of serum metabolites. The metabolite features and AUC curves for random forest analysis of COVID-19 patients with mild/moderate disease compared to those with severe disease (a); COVID-19 patients with a fatal outcome compared to those with mild/moderate disease (b); COVID-19 patients that survive following severe disease compared to those that don’t survive following severe disease (c).

**Supplementary Fig. 4.**
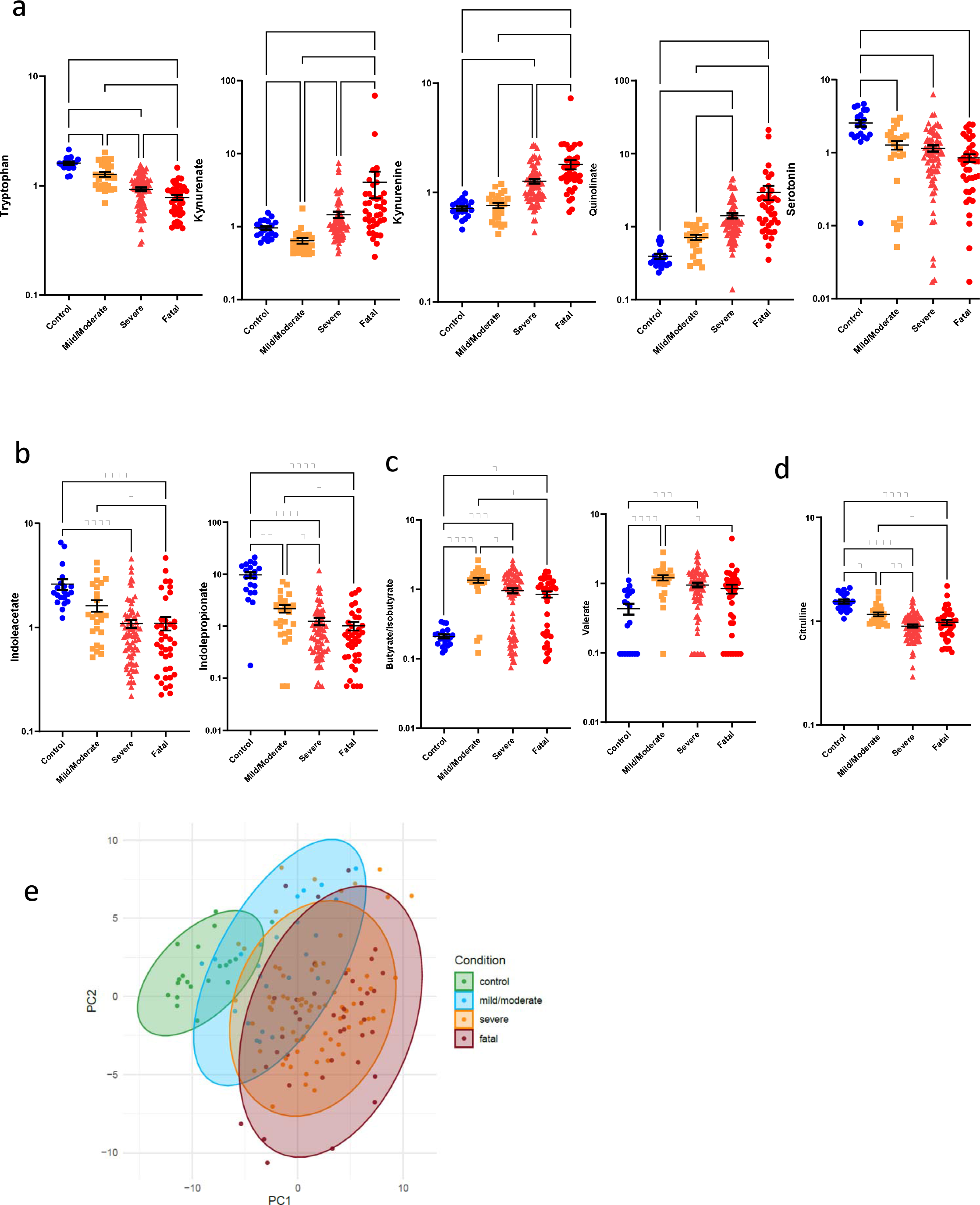
Serum microbial metabolites. Representative examples of metabolites generated by host metabolism of tryptophan (a). Selected examples of serum levels of microbial metabolites due to tryptophan metabolism (b) or SCFAs (c). Serum citrulline levels (d). PCA plot illustrates the differences in serum metabolites associated with microbial metabolism (e). Differences between groups are calculated using the Kruskal-Wallis test and Dunn’s multiple comparison test (*p<0.05, **p<0.01, ****p<0.0001).

**Supplementary Fig. 5.**
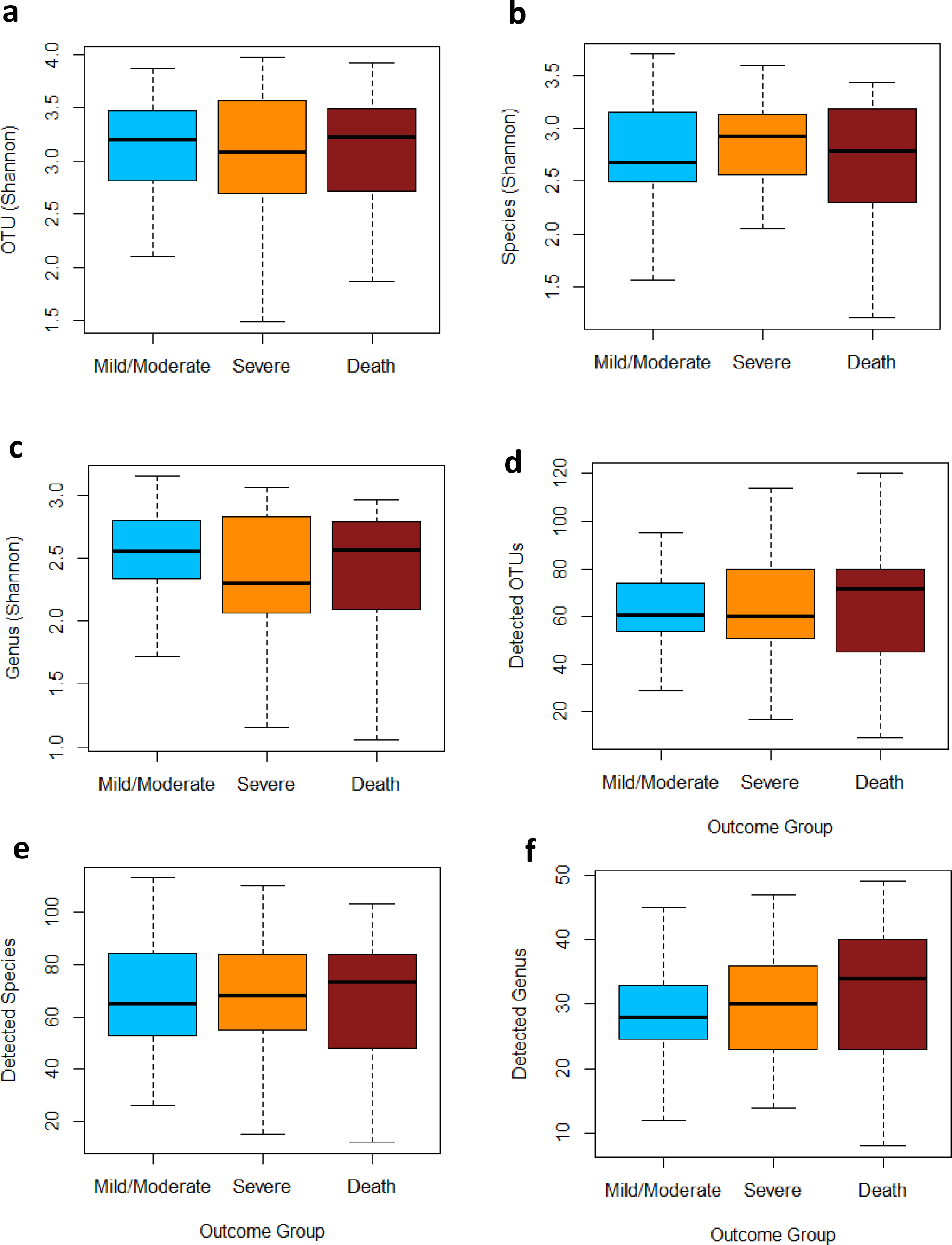
Gut microbiome alpha diversity. Boxplots showing the variation of the Shannon Diversity and Detected taxa for the gut microbiome profiles for the three outcome groups at OTU (a and d), Species (b and e) and Genus (c and f) levels.

**Supplementary Fig. 6.**
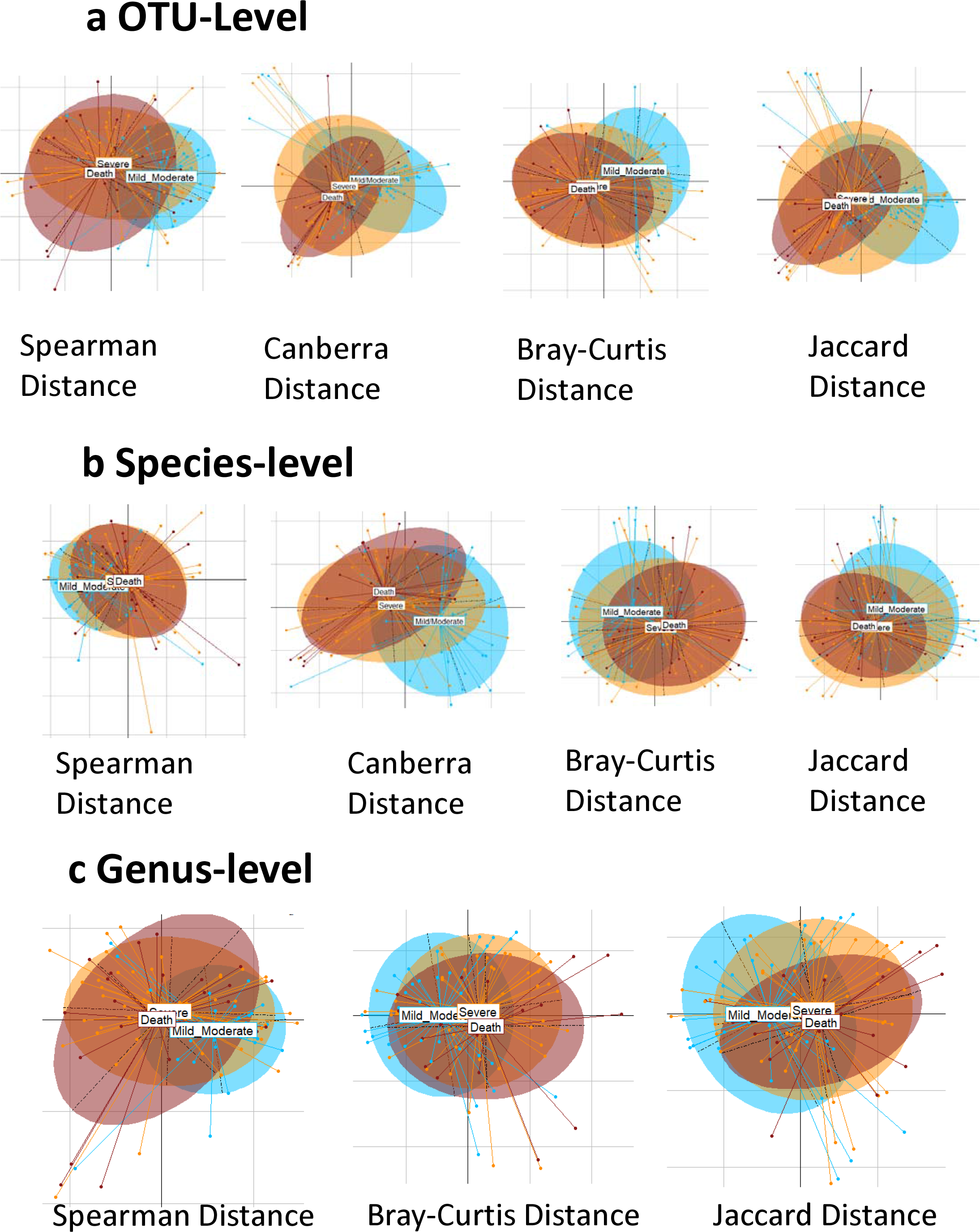
Gut microbiome beta diversity. Principal coordinate analysis showing the resolution of the gut microbiome profiles from the 99 patients belonging to the three outcome groups at (a) OTU and (b) Species level, obtained using four different distance measures. (c) Principal coordinate analysis showing the resolution of the gut microbiome profiles from the 99 patients belonging to the three outcome groups at the genus level obtained using the Spearman, Bray-Curtis and Jaccard distance measures.

**Supplementary Fig. 7.**
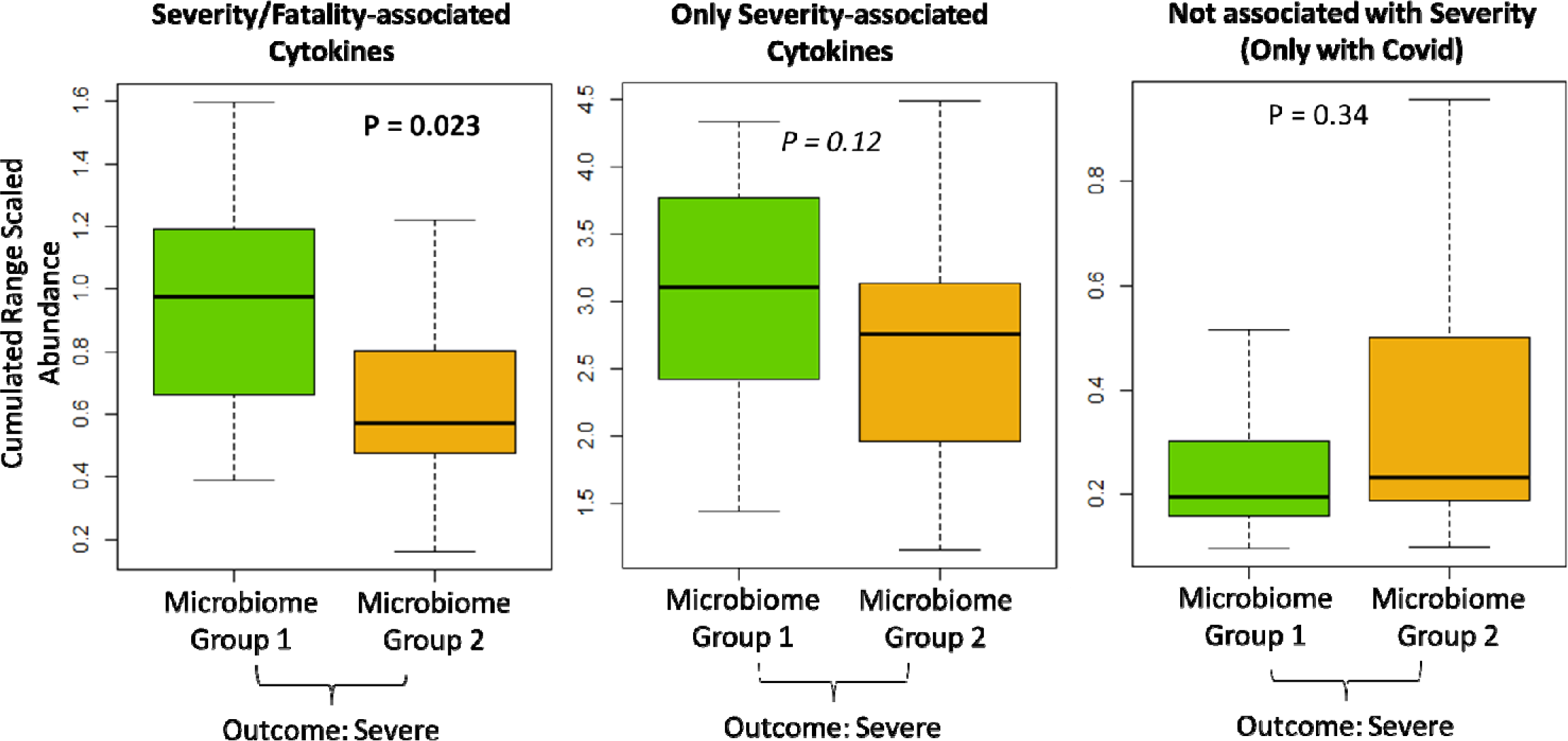
Cytokine levels associated with Microbiome Groups. Boxplot showing the differences in the cumulated range-scaled levels of the three groups of elevated cytokines between surviving patients with severe symptoms who had a high-risk MicrobiomeGroup1 and those patients with severe symptoms who were classified to the low-risk MicrobiomeGroup2. The p-values of the Mann-Whitney tests obtained for the comparisons within the three groups of cytokines are indicated. Each cytokine level was range-scaled across patients to a value between 0 and 1. For each patient, the range-scaled values of all cytokines within the same group were then cumulated by adding the corresponding range-scaled values obtained for the given patient.

**Supplementary Fig. 8.**
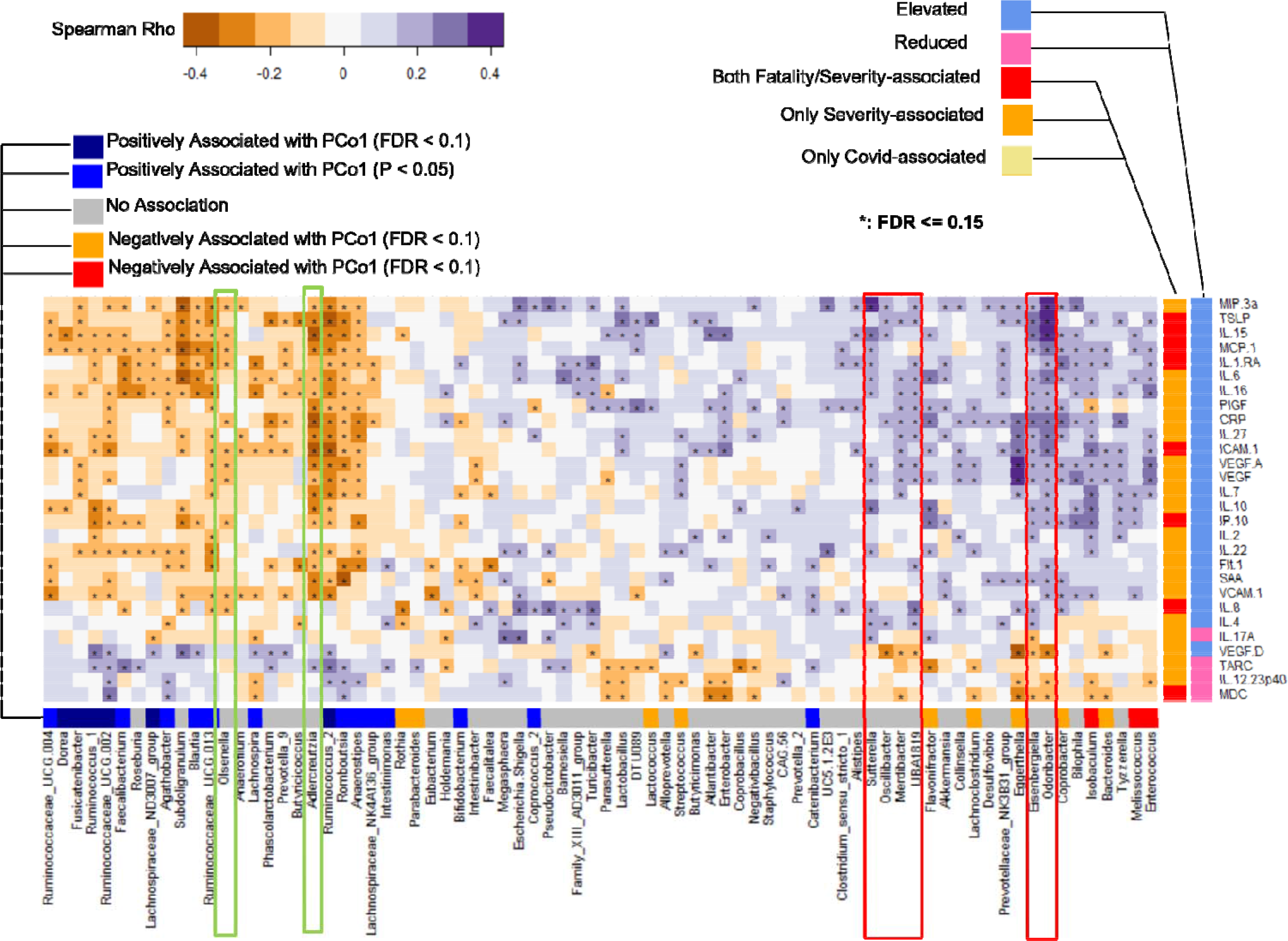
Bacterial genera correlate with circulating inflammatory mediators. Heatmap showing the Spearman correlations between the 73 genus-level markers (detected in at least 5 of the 99 patients and showing association with FDR ≤ 0.15 with at least one of the severity/fatality-associated cytokines) and 28 severity/fatality-associated cytokines. The groups of the different cytokines and their direction of change (elevated or reduced) with severity/fatality are also indicated in specific colors. Also indicated are the patterns of the various genus-level markers with PCo1 that resolves the two microbiome groups (positive PCo1 with MicrobiomeGroup2). The genera whose associations are indicated in red boxes are those that did not show any associations with either of the two Microbiome configurations (or groups) in terms of their association with PCo1, but independently show association with an inflamed host phenotype. The genera in green boxes show the opposite trends with the inflamed phenotype.

**Supplementary Figure 9.**
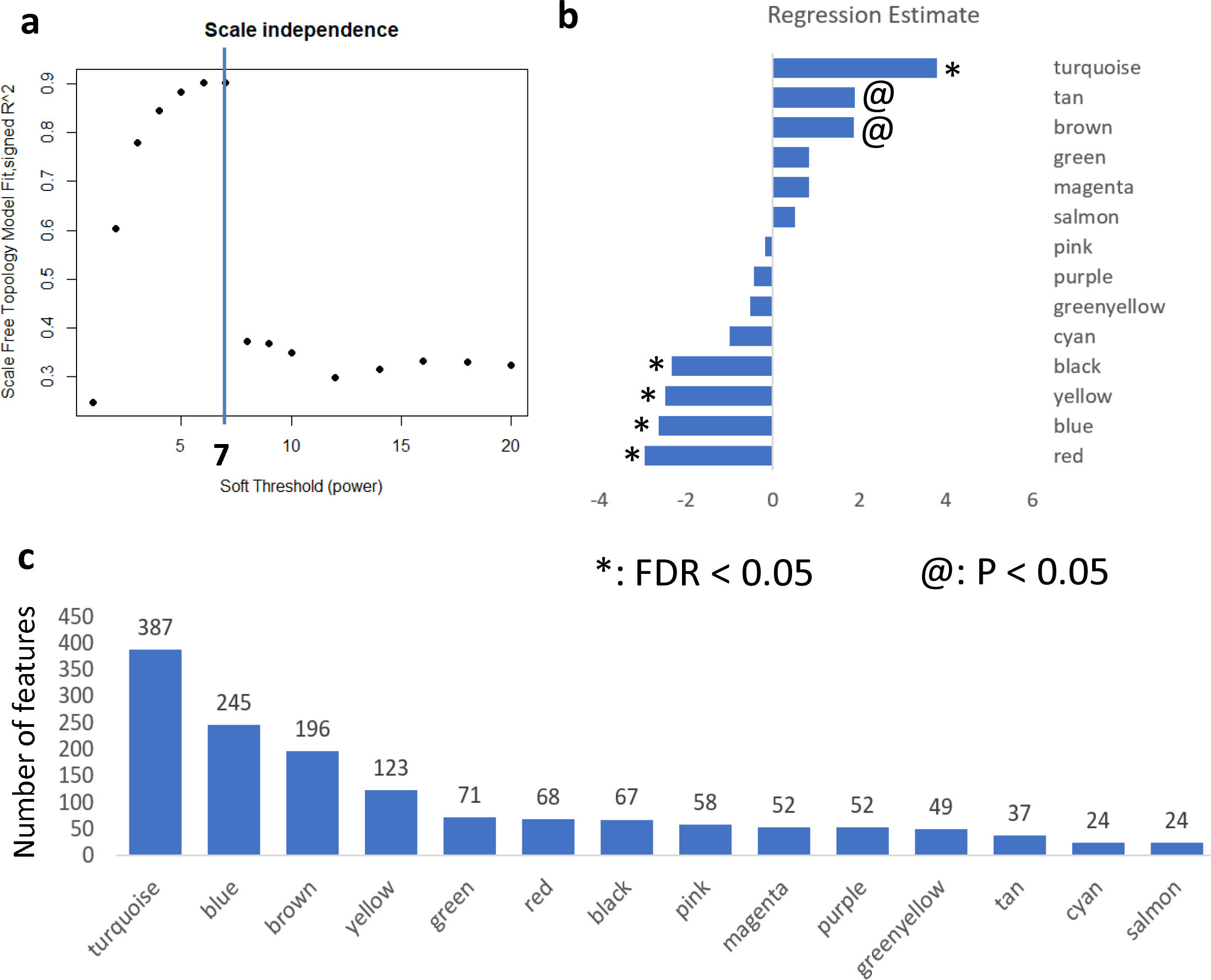
Overview of the steps and the results of combined WGCNA from the three data layers. (A) shows the Scaled-Independence plot and the Scale-free topology fit and highlights the selection of the soft-power of 7 as it has the maximum scale-free nature for the network. (B) Shows the regression coefficients of the 14 modules obtained using Ordinary Least-square Regression for worse outcome (where in the outcomes were ranked as 1 for mild and moderate; 2 for severe and 3 for death). The modules with significant (Benjamini-Hochberg corrected FDR ≤ 0.05) and nominal associations (P ≤ 0.05) are also indicated. (C) Shows the sizes of the different modules in terms of the feature.

**Supplementary Fig. 10.**
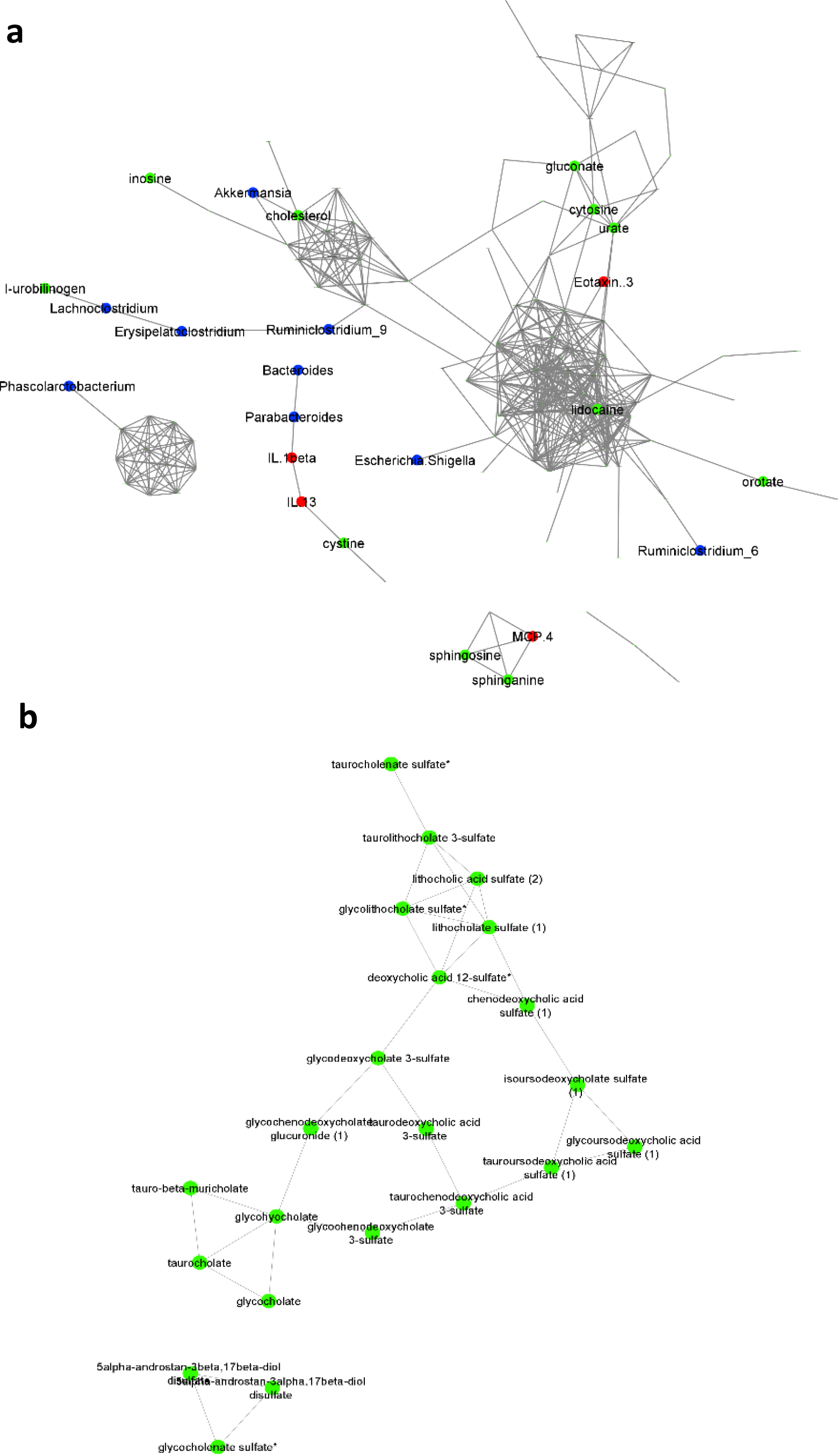
Modules showing nominal positive associations with severity and death. Positive association networks obtained for the features affiliated to (a) brown and (b) the tan module, using the ccrepe approach (Spearman correlation, iterations = 1000, p ≤ 0.01). Key taxa and metabolites are highlighted.

**Supplementary Fig. 11.**
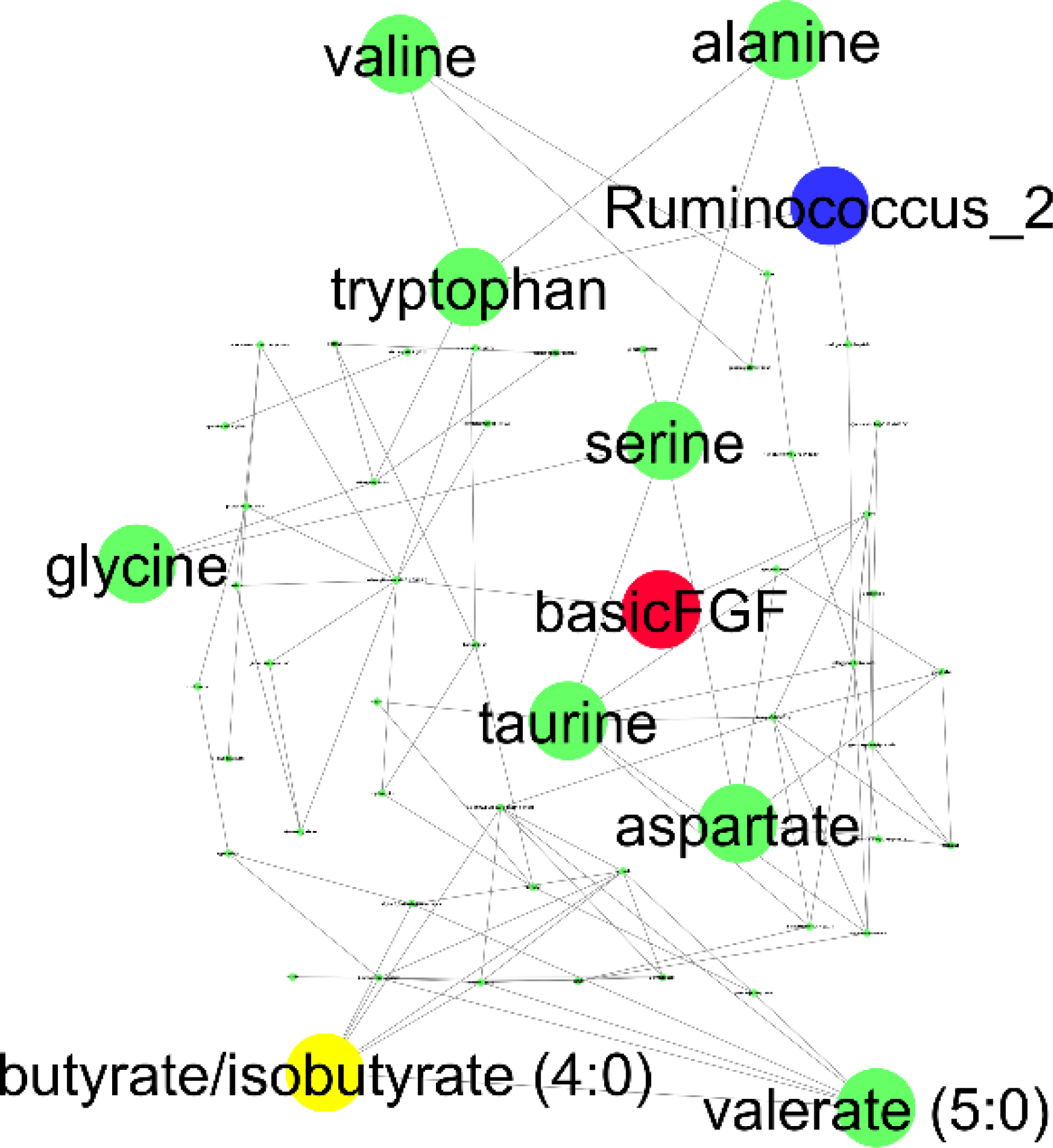
Association patterns within the ‘red’ module. Positive association networks obtained for the features affiliated to the red module, using the ccrepe approach (Spearman correlation, iterations = 1000, p <= 0.01). Key taxa and metabolites are highlighted.

**Supplementary Fig. 12.**
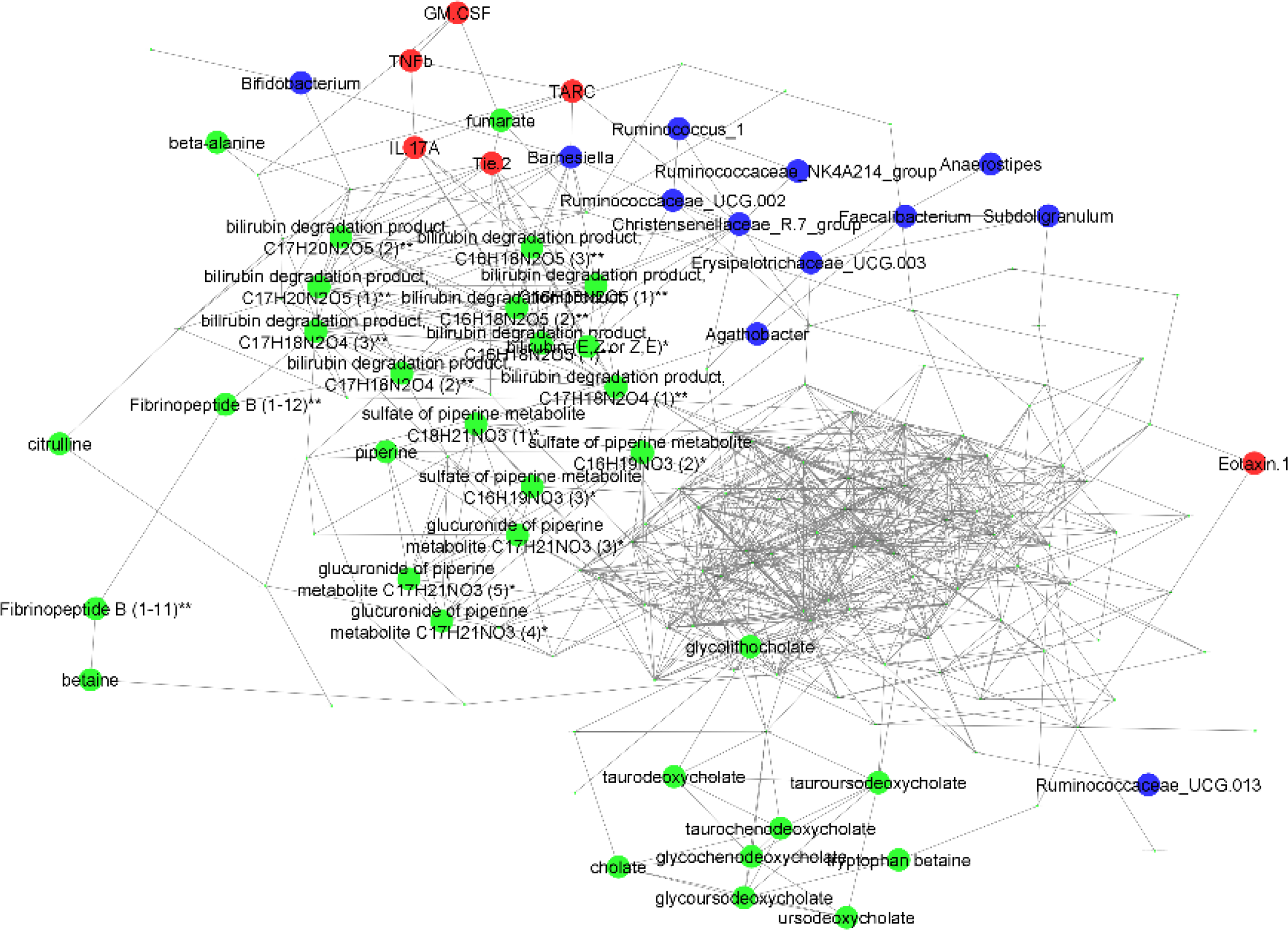
Association patterns within the ‘blue module. Positive association networks obtained for the features affiliated to the blue module, using the ccrepe approach (Spearman correlation, iterations = 1000, p <= 0.01). Key taxa and metabolites are highlighted.

**Supplementary Fig. 13.**
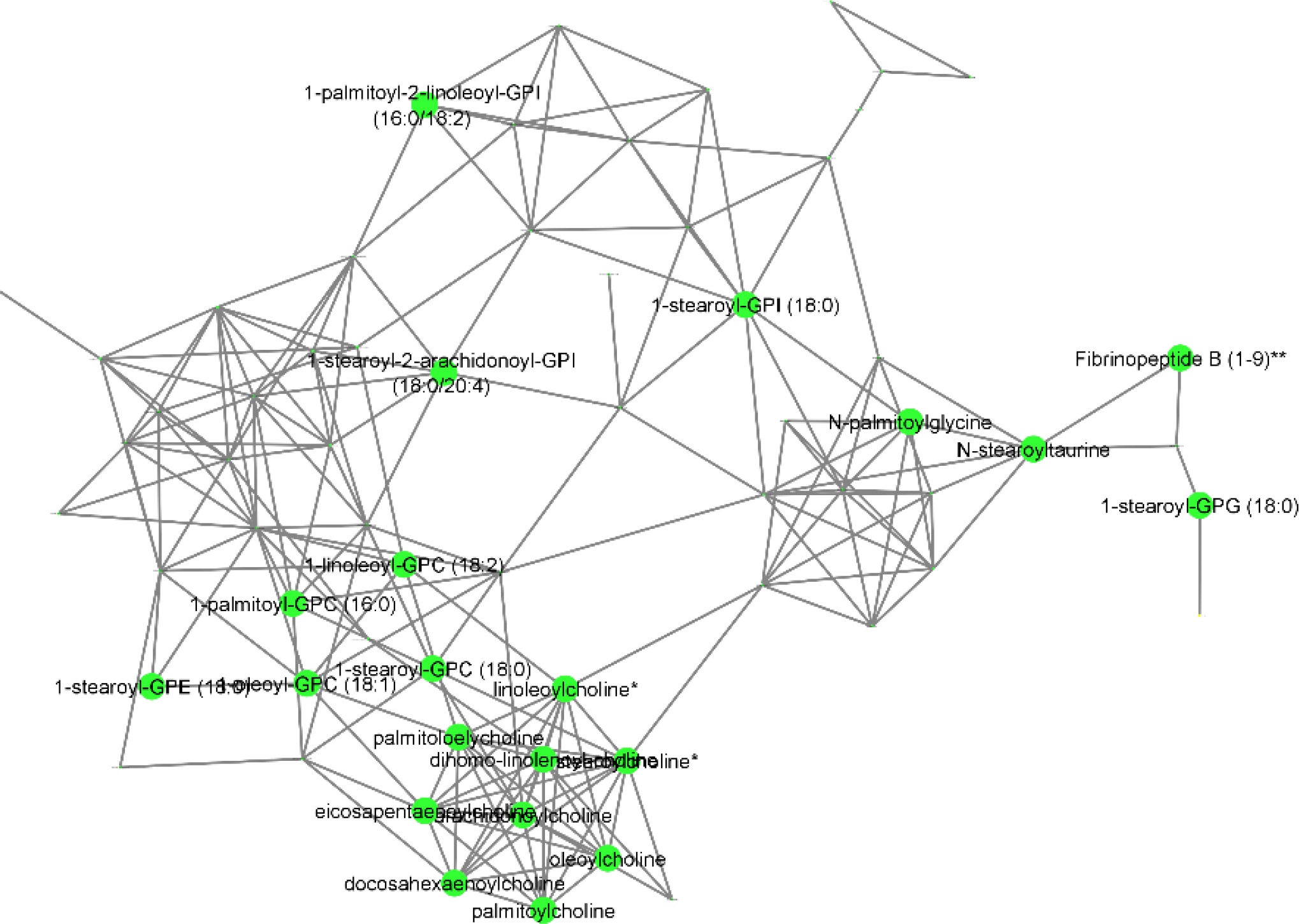
Association patterns within the ‘black’ module. Positive association networks obtained for the features affiliated to the black module, using the ccrepe approach (Spearman correlation, iterations = 1000, p <= 0.01). Key taxa and metabolites are highlighted.

**Supplementary Fig. 14.**
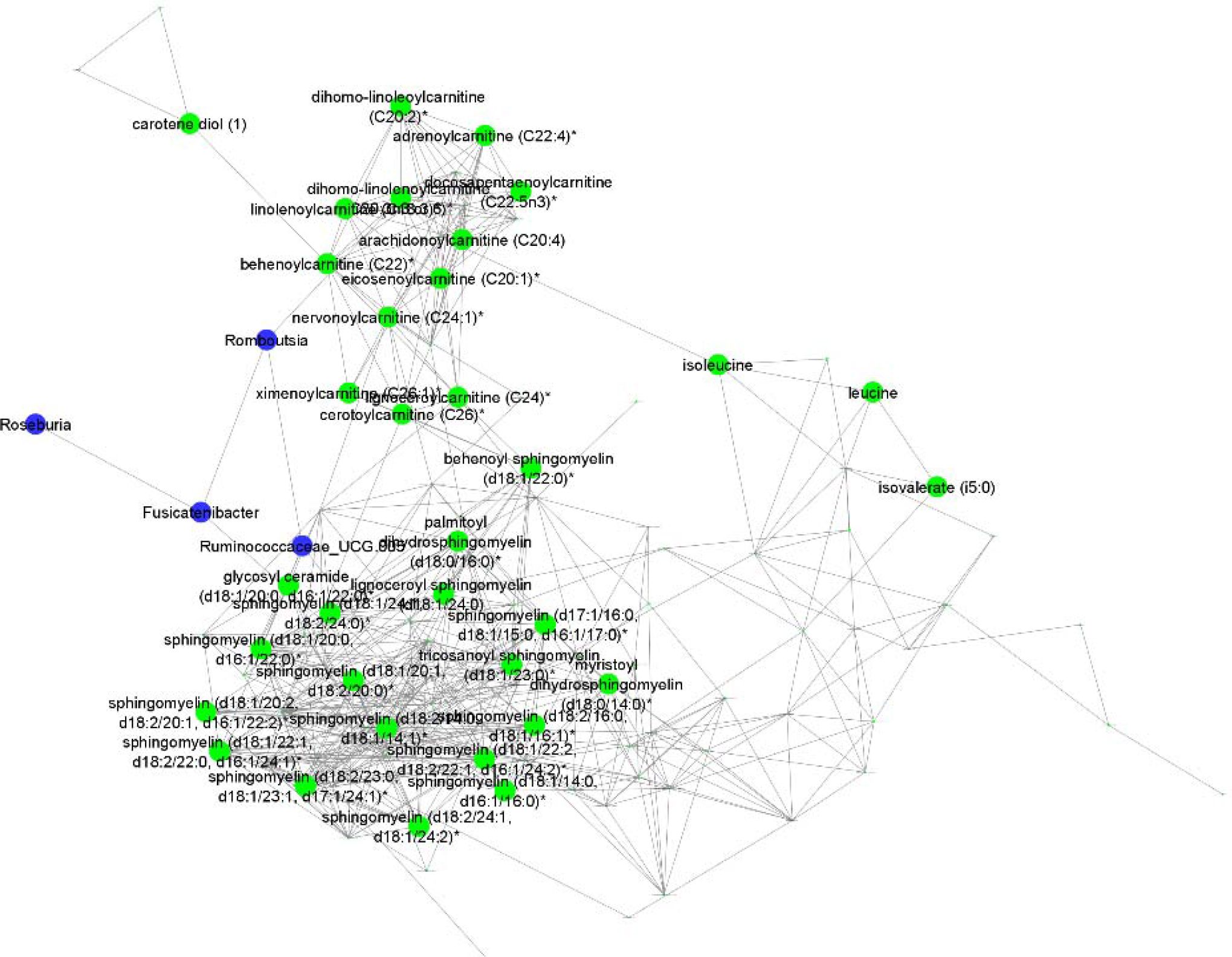
Association patterns within the ‘yellow module. Positive association networks obtained for the features affiliated to the yellow module, using the ccrepe approach (Spearman correlation, iterations = 1000, p <= 0.01). Key taxa and metabolites are highlighted.

